# Animal Emotions and Consciousness: Researchers’ Perceptions, Biases, and Prospects for Future Progress

**DOI:** 10.1101/2023.10.12.562087

**Authors:** Matthew N Zipple, Caleb Hazelwood, Mackenzie F Webster, Marcela E Benítez

## Abstract

Do animals have emotions? Scientists and philosophers have long struggled with this question, with debates ranging from whether animals experience an “internal world” to whether we are capable of studying it. Recently, theoretical, and methodological advances have rekindled this debate, yet, it is unclear where the scientific consensus on these topics lies today. To address this gap, we administered a survey of professional animal behavior researchers to assess perceptions regarding (1) the taxonomic distribution of emotions and consciousness in non-human animals, (2) respondents’ confidence in this assessment, and (3) attitudes towards potential for progress and possible pitfalls when addressing these questions. In general, animal behavior researchers (n=100) ascribed emotionality and consciousness to a broad swath of the animal taxa, including non-human primates, other mammals, birds, and cephalopods, with varying degrees of confidence. There was a strong positive relationship between how likely a respondent was to attribute emotions to a given taxa and their confidence in that assessment, with respondents assuming an absence of emotions and consciousness when they were unsure. In addition, respondents’ assessments were shaped by several traits (e.g., advanced cognitive abilities, consciousness) that they also admitted were not necessary for an animal to experience emotions. Ultimately, a large majority of researchers were optimistic that tools either currently exist or will exist in the future to rigorously address these questions (>85%) and that animal behavior, as a field, should do more to encourage emotions research (71%). We discuss implications of our findings for publication bias, ethical considerations, and identify an emergent consensus for the need of a functional definition of emotions to facilitate future work.

**Significance Statement:** Emotions and consciousness are fundamental components of human experience—these phenomena are central to our behavior, relationships, and sense of meaning. Whether these experiences are shared by non-human animals has long been a subject of philosophical and scientific debate. In this paper we describe, for the first time, results from a survey of animal behavior researchers regarding their perceptions of these questions and the ability of science to answer them. Researchers ascribe emotions and consciousness to many taxa, and their likelihood of doing was strongly predicted by phylogeny and researchers’ confidence in their answers. We hope these results spur additional interdisciplinary collaboration to rigorously pursue these questions and create a baseline for future comparisons to track scientific attitudes over time.

> “I have chosen bats instead of wasps or flounders because if one travels too far down the phylogenetic tree, people gradually shed their faith that there is experience there at all”--Thomas Nagel

## Introduction

Do animals experience emotions? For centuries, this seemingly simple question has been debated by philosophers and scientists, with some suggesting that emotions are uniquely human traits and others arguing that emotions play a significant role in how animals behave (1–4). For example, Aristotle and Hume believed that animals and humans share similar desires and emotions, with Hume going so far as to extend thought and reason to non-human animals (1, 3). On the other hand, the French philosopher Descartes challenged these beliefs, arguing that animals were mere machines, lacking the capacity for emotions or consciousness (2). The American philosopher Thomas Nagel took a middle path, arguing that although many animals clearly have their own subjective experiences, those emotions and mental states “may be permanently denied to us by the limits of our nature” and therefore make poor subjects of scientific study (4).

In the 19^th^ and 20^th^ century, scientific discourse on the question of animal emotions paralleled this philosophical dispute, with evolutionary biologists, behaviorists, and ethologists debating both the presence of emotions in animals and our ability as scientists to critically evaluate a fundamentally “private phenomenon” (5). Darwin displayed no compunction about ascribing emotions to animals, arguing in *The Expression of the Emotions in Man and Animals* that animals display and express many of the same emotions that humans do (6). In *The Descent of Man, and Selection in Relation to Sex*, Darwin claimed that “the lower animals, like man, manifestly feel pleasure and pain, happiness, and misery” (7). But within 50 years of Darwin’s work, behaviorist theories had come to dominate animal behavior research. John B. Watson and B.F. Skinner, among others, frowned at the notion of studying animal (and human) emotions because they considered it unscientific. For behaviorists, only behavior—specifically behavior as a response to external stimuli—“counted” as observable and quantifiable data. They argued that even if subjective experiences, like emotions, did exist in non-human animals, emotions made poor study questions because they were unobservable, unmeasurable, and unverifiable (8). This sentiment can perhaps be best summed up by the evolutionary biologist George Williams (1992) who wrote, “I am inclined merely to delete it [the mental realm] from biological explanation, because it is an entirely private phenomenon, and biology must deal with the publicly demonstrable” (5).

One of the issues that plagues all researchers of emotion, and animal emotion researchers in particular, is the inconsistency in definitions across fields. Indeed, one argument for why our understanding of emotions has lagged behind other areas of cognition is that there remains fundamental disagreement on the definition of emotion (9). This hurdle is particularly large for animal researchers, because many definitions of emotions include the subjective aspect as a fundamental part of emotional experience – something that is difficult, if not impossible, to operationalize in a non-human animal. Frans de Waal neatly summarized the dilemma animal researchers face, noting that while emotions in animals are rarely denied, their importance is often questioned leaving us “with the curious situation that a widely recognized aspect of animal behavior is deliberately ignored or minimized” (10).

Despite these hurdles, research in recent decades demonstrates that, at minimum, animals experience changes in their core affective states, categorized by a spectrum of valence and arousal levels (11, 12). These conclusions come from advances in the fields of psychology, ethology, animal cognition, neuroscience, and behavioral endocrinology, that have allowed researchers to monitor and examine internal states in animals. For example, comparative psychologists and neuroscientists have continued Darwin’s approach to studying animal facial expressions, now with the added scientific power of machine learning and neurobiological recordings (13, 14). Neuroscientists have focused on identifying the complex relationship between emotions and neural structures of the brain, highlighting several areas of the cerebral cortex and subcortical structures (e.g., amygdala) that can initiate and regulate emotions. These regions appear homologous in other animal species, suggesting that they likely play a similar role (reviewed in (15)). These discoveries and others have opened the door for behavioral ecologists to explicitly incorporate animal emotions into theories of why and how animals do what they do. For example, Crump et al. (2020) argue that animal emotions and mood play a critical role in explaining winner and loser effects in animal contests (16).

The distribution of opinions regarding the presence and importance of animal emotions has fluctuated substantially over time, so where are we now? In this paper, we set out to quantify the current distribution of expert thought on these topics by surveying animal behavior researchers across different fields and academic stages. The main goals of this study are to quantify researchers’ current positions on animal emotions in general (“do animals have emotions? how confident are you in that assessment?”), the taxonomic distribution of animal emotions (“which animals have emotions?”), and whether these questions are answerable (“do we have the tools to study animal emotions?”). By making clear the current distribution of thought, we hope to instigate interdisciplinary conversation and help guide future research initiatives in this growing field.

When we say that animals do or do not have emotions, we are making a philosophical claim that depends on the operative concept of “emotion” we have in mind. Different researchers have different criteria for what emotions are in the first place. In such circumstances, researchers studying the exact same model system and working with the exact same empirical data may produce opposite verdicts about the presence and importance of animal emotions. To investigate this philosophical question, we rely on a methodology referred to as “experimental philosophy”. Experimental philosophy combines the conceptual frameworks traditionally associated with philosophy and the experimental rigor of cognitive science (17). It uses survey data to generate understanding about the concepts that scientists use— concepts that are instrumental for their research—instead of legislating those concepts from the armchair. Similar methods have recently been used to examine differences in how geneticists define genes (18–20), as well as biologists’ positions on “innateness” (21, 22) and evolutionary and ecological models (23). Thus, by applying an empirical approach to a philosophical concept such as “emotion,” we are better poised to appreciate who, among animal researchers, ascribe emotions to their study system; why they do or do not believe such ascriptions are warranted; when, and under what conditions, such ascriptions are warranted; and what factors (such as their beliefs about consciousness, concerns about anthropomorphism, etc.) are predictors of certain beliefs.

To our knowledge there have been no previous attempts to quantify the distribution of thought regarding animal behavior researchers’ perceptions of the taxonomic distribution of animal emotionality. Here we take the first step towards filling this gap by conducting a survey of professional animal behavior researchers, asking them to describe their beliefs about the taxonomic distribution of animal emotions and consciousness and their confidence in those assessments. The result is the first assessment of how animal behavior researchers, as a group, think about the importance of emotions and consciousness in non-human animals.

## Methods

In this study, we collected survey data from professional animal behavior researchers who provided responses to multiple-choice questions, free-form text fields, and Likert scales (24). Responses were solicited through two primary means. First, we emailed all departments on U.S. News and World Reports’ list of best graduate school programs in behavioral neuroscience, cognitive psychology, ecology/evolutionary biology, and neuroscience/neurobiology, requesting that our survey be distributed to relevant members of their department. Second, we posted solicitations of our survey on Twitter (now X), explaining the type of respondents that we were seeking and providing a basic overview of the types of questions respondents would be asked (See Supplemental Materials for full text of solicitations).

Respondents first provided individual data about their career stages, research disciplines, study systems, and interactions with domesticated animals. They then answered a series of questions about whether they ascribe emotional responses and consciousness to various taxa, as well as the relevance of various factors (such as phylogenetic proximity to humans, social behavior, group living, facial expressions, etc.) to their decisions, and their confidence in their assessments. Furthermore, we surveyed the participants’ meta-level beliefs about the role of emotions in their discipline: Does talk of animal emotions risk anthropic fallacies? Should research into non-human animal emotions be encouraged? Does the participant feel comfortable voicing their opinions about non-human animal emotions in professional settings? We have included the full survey in the Supplementary Materials.

### Inclusion criteria

To guarantee that respondents were academic researchers and had completed the survey with care, we included only respondents who (1) identified as professional scientists or current academic students and (2) navigated all the way to the end of the survey. As a basic test of an individual carefully considering the questions and giving good-faith responses, we included only respondents that (3) identified either “all or nearly all” or “most” humans as displaying emotions.

### Statistical analyses

We performed all statistical analyses in R (version 4.2.2, (25)). For mixed effects models we used the package glmmTMB (version 1.1.5, (26)). We created Likert scale visualizations using the likert package (version 1.3.5, (27)). Details on model structure, including random and fixed effects are provided in the results.

### Ethical Approval

Because we did not store sensitive personal information and our survey did not pose a risk of adverse outcomes for participants, we obtained institutional review board exemptions from each of our institutions (Cornell University, Duke University, and Emory University).

## Results

### Respondent Characteristics

In total, we received 100 complete responses to our survey that met the inclusion criteria. 45% of respondents were graduate students, 31% were faculty, and 20% were post docs (the remainder fell into other professional categories, see Table 1). Most respondents identified as belonging to multiple fields and most studied more than one taxon. Our sample included a large number of behavioral ecologists (63%), evolutionary biologists (38%), and neurobiologists (26%), as well as a smaller number of biological anthropologists (13%), biological psychologists (12%), and cognitive psychologists (11%). The most common taxa of study among respondents were birds (43%), non-human primates (32%), and other mammals (48%), though each of the taxa that we asked respondents to assess were studied by at least some members of our sample (Table 1).

**Table 1.** Academic traits of included respondents to our survey.

|  |  | Percentage of Respondents (n =100) |
| --- | --- | --- |
| ***Career Stage*** |  |  |
|  | Graduate Student | 45 |
|  | Postdoctoral Researcher | 20 |
|  | Assistant Professor or Equivalent | 13 |
|  | Associate Professor or Equivalent | 10 |
|  | Full Professor or Equivalent | 5 |
|  | Emerita | 2 |
|  | Undergraduate Student | 2 |
|  | Other PhD Researcher or Adjunct Faculty | 3 |
| ***Self-Identified Fields of Study*** |  |  |
|  | Behavioral Ecologist | 63 |
|  | Evolutionary Biologist | 38 |
|  | Biological Anthropologist | 13 |
|  | Neuroscience | 26 |
|  | Cognitive Psychology | 11 |
|  | Biological Psychology | 12 |
|  | Other | 26 |
| ***Self-Identified Taxa of Study*** |  |  |
|  | Non-human Primates | 32 |
|  | Other Mammals | 48 |
|  | Birds | 43 |
|  | Amphibians or Reptiles | 11 |
|  | Fish | 15 |
|  | Squids, Octopus, or Cuttlefish | 2 |
|  | Insects | 16 |
|  | Other Invertebrates | 8 |

### Perceived taxonomic distribution of emotions in non-human animals

Overall, animal behavior researchers identify a wide taxonomic breadth of animals as experiencing emotions that shape their behavior. Majorities of those surveyed ascribed emotions to “most” or “all or nearly all” non-human primates (98%), other mammals (89%), birds (78%), octopus, squids, and cuttlefish (72%), and fish (53%). And for all taxa considered, a majority of researchers ascribe emotions to at least some members of each taxon, including insects (67%) and other invertebrates (71%). We present our full results in Figure 1.

**Figure 1.**
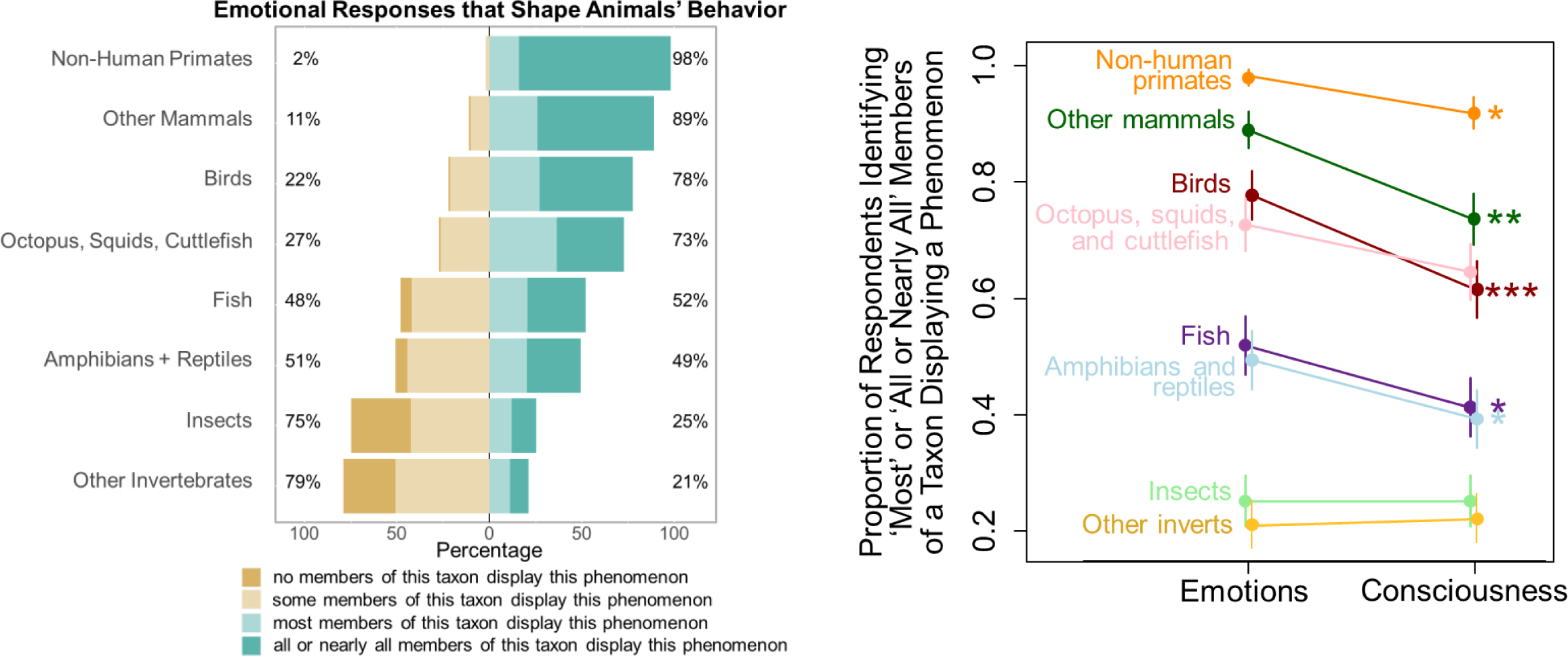
(Left) Respondents’ assessment of the distribution of emotional responses that shape animals’ behavior across a range of non-human taxa. (Right) For most taxa assessed, respondents were more likely to ascribe emotional responses to animals than they were consciousness. Asterisks represent significance: * p <0.05; ** p < 0.01; *** p < 0.001.

### Perceived taxonomic distribution of consciousness in non-human animals

Animal behavior researchers also ascribed consciousness to a broad taxonomic breadth of animals, although a more restricted range as compared to emotions. As with emotions, majorities ascribed consciousness to “most” or “all or nearly all” non-human primates (92%), other mammals (73%), octopuses, squids, and cuttlefish (64%), and birds (61%). As with emotions, most researchers ascribe consciousness (Figure S1) to at least some members of each taxon, including insects (51%) and other invertebrates (52%).

Each respondent assessed the frequency of emotions and consciousness across multiple taxa. Combining each of these responses into a mixed effects model with random effects of respondent ID and taxon ID revealed that respondents were substantially more likely to ascribe emotions to most or all members of a taxa than they were to ascribe consciousness to the same (p < 0.0001). When breaking these differences down within different taxa (Figure 1), respondents were more likely to ascribe emotions than consciousness to non-human primates (p = 0.047), other mammals (p = 0.002), birds (p = 0.0009), fish (p = 0.02), and amphibians and reptiles (p = 0.04)

### Respondents’ confidence in their assessments of animals’ emotions and consciousness

Respondents displayed a wide range of confidence in their assessment of the distribution of emotions and consciousness, depending on the taxon in question (Figure 2). Respondents were most confident in their assessments of non-human primates (mean = 4.5) and other mammals (mean = 4.2), followed by birds (mean = 3.7). Respondents’ confidence declined when considering ectotherms, with respondents less confident about their assessment of cephalopods (mean = 3.2), followed by fish (mean = 2.9), amphibians and reptiles (2.8), insects (2.5), and finally other invertebrates (mean = 2.4). Confidence regarding assessments of consciousness followed an identical taxonomic pattern as did confidence in assessment of emotions. Generally, confidence in assessment of the two phenomena were quite similar (note mostly overlapping 95% confidence intervals in Figure S2).

**Figure 2.**
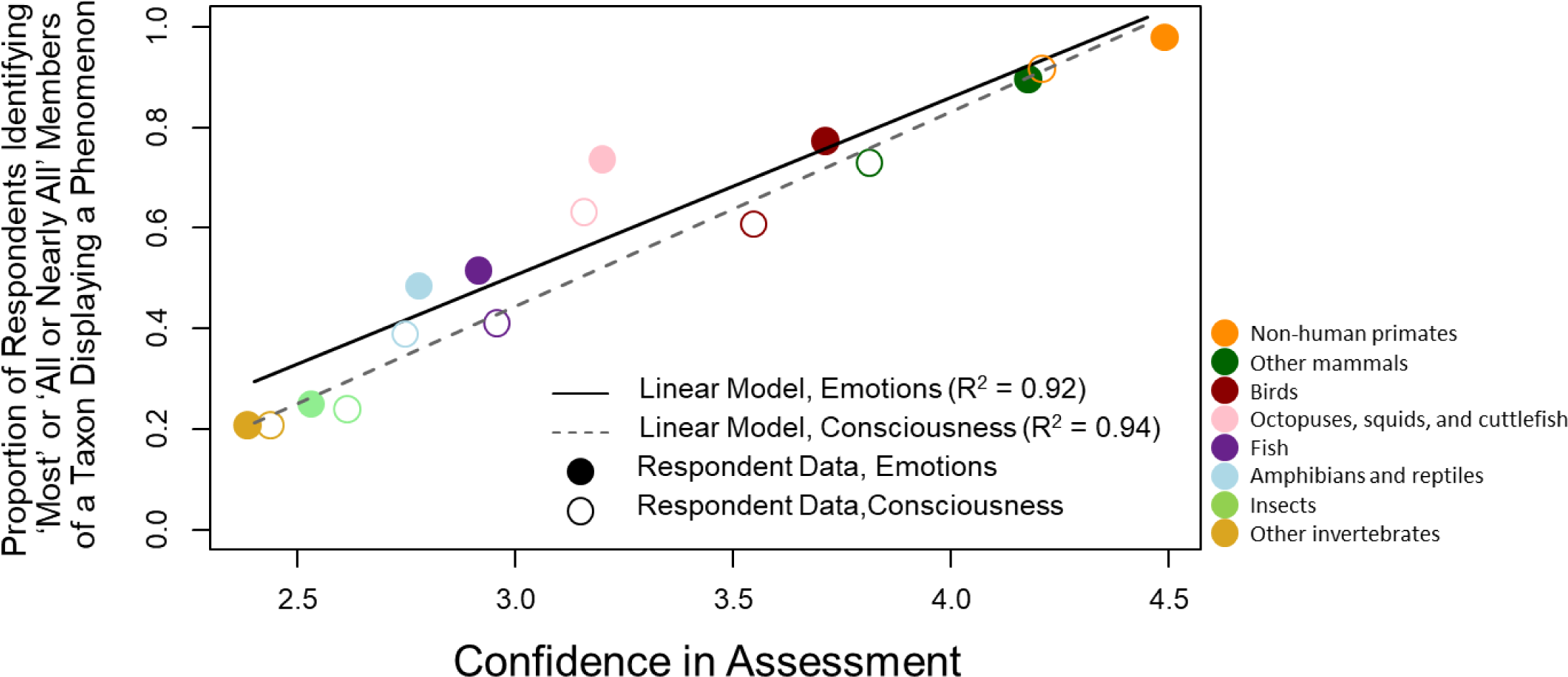
Respondents’ assessment of the distribution of emotions or consciousness in a given taxon was almost perfectly predicted by how confident they were in that assessment. For each taxon, respondents provided their confidence in their assessment 1 (“not at all confident”) to 5 (“certain”). For both assessments of emotions (R^2^=0.92) and consciousness (R^2^=0.94), respondents ascribed phenomena more broadly within a taxon when they were more confident in their assessment.

The concordance between respondents’ assessment of the distribution of emotions and consciousness in a given taxon and *their confidence* in that same assessment is striking—as a group, respondents’ average confidence in their assessment of emotionality for a given taxon almost perfectly predicted the distribution of emotions of that they ascribed to that taxon (R^2^ = 0.92, p = 0.001, Figure 2).

### Additional biases shaping researchers’ assessments of animal emotions

To examine potential biases shaping responses, we asked respondents to think about their responses and assess how important various animal characteristics were in shaping those assessments (“not at all important”, “somewhat important”, “very important”). We then asked respondents to assess the extent to which those same characteristics were necessary for an animal to experience emotions (“not at all necessary,” “somewhat necessary”, “essential”). Thus, respondents were given the opportunity to provide answers that were logically at odds with each other, indicating a bias in scientists’ thinking on these topics. Specifically, respondents could identify a characteristic as being “not at all necessary” for an animal to experience emotions, while still identifying these same characteristics as “somewhat” or “very” important to their answers.

For nearly every characteristic that we asked about, we detected a discrepancy between what respondents identified as important (‘somewhat’ or ‘very’) in shaping their assessments of whether animals experience emotions and what they identified as necessary (‘somewhat necessary’ or ‘essential’) for animals to experience emotions (Figure 3). Remarkably, two characteristics (facial expressions and phylogenetic closeness to humans) were identified by large majorities of respondents as not at all necessary (91% and 81% respectively) but were also identified by majorities (51% and 68%) as at least somewhat important in shaping their assessments. Sizable minorities also identified domestication (25%) and use of language (44%) as important in shaping their assessments even though many fewer identified these same characteristics as at all necessary for animals to experience emotions (6% and 15% respectively). Even when large portions or majorities of respondents identified a given characteristic as necessary for animals to experience emotions, even more respondents identified these characteristics as important in shaping their thinking, including group living (34% somewhat necessary or essential vs 60% somewhat or very important), being highly social (46% vs 67%), and advanced cognitive abilities (64% vs 87%).

**Figure 3.**
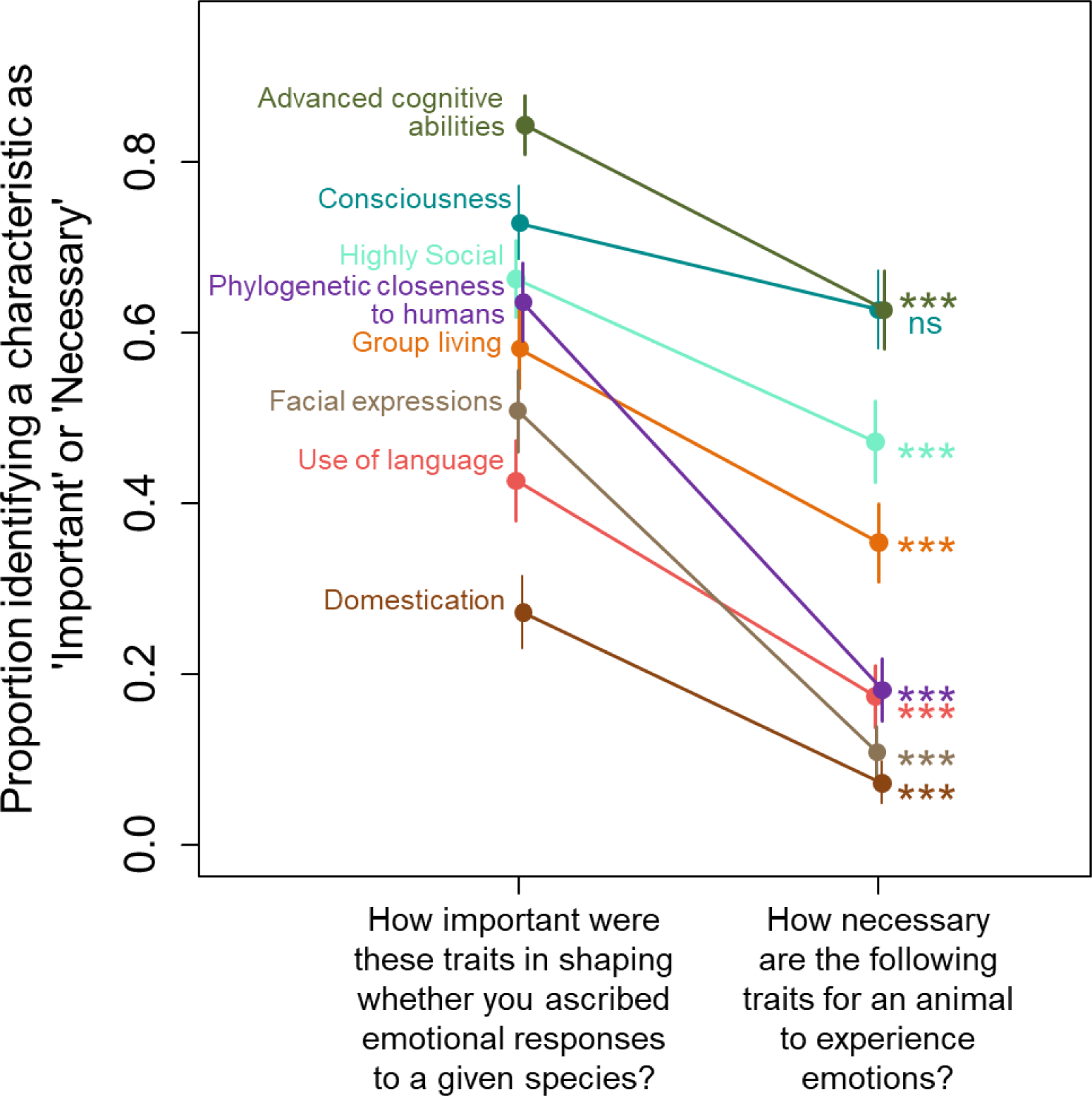
A mismatch exists in respondents’ sense of whether a given trait is necessary for animals to experience emotion (right) and their own assessment of how important that same characteristic was in shaping whether they ascribed emotional responses to a given species (left). Respondents’ assessments were shaped by a number of traits that they admitted were not necessary for an animal to experience emotions. Asterisks represent significance: *** p < 0.001.

The one characteristic that respondents identified as both important and necessary without a significant difference (p > 0.05) in their assessment was consciousness. 75% of respondents identified animals’ consciousness as important in shaping their view of whether those animals had emotions, and 65% said that consciousness was at least somewhat necessary for animals to experience emotion. This latter pattern mirrors the tight perceived taxonomic linkages between emotions and consciousness that respondents had previously expressed (Figure 1).

### Perceptions of the risk of anthropic fallacies

Respondents were divided on the topic of the risks posed by the twin anthropic fallacies of anthropomorphism and anthropodenial (28, 29). Overall, 49% agreed that “Discussing the role of emotions in non-human animals’ behavior risks inaccurately projecting human experience onto study subjects (i.e. anthropomorphizing),” while 46% disagreed (5% neither agreed nor disagreed). Respondents had a more universal view regarding anthropodenial, with 89% of respondents agreeing that “Failing to consider the role of emotions in non-human animals’ behavior risks ignoring homologous mechanisms that are involved in both human and non-human behavior (i.e. anthropodenial)” and only 6% disagreeing (5% neither agreed nor disagreed). Full results are displayed in Figure 4.

**Figure 4.**
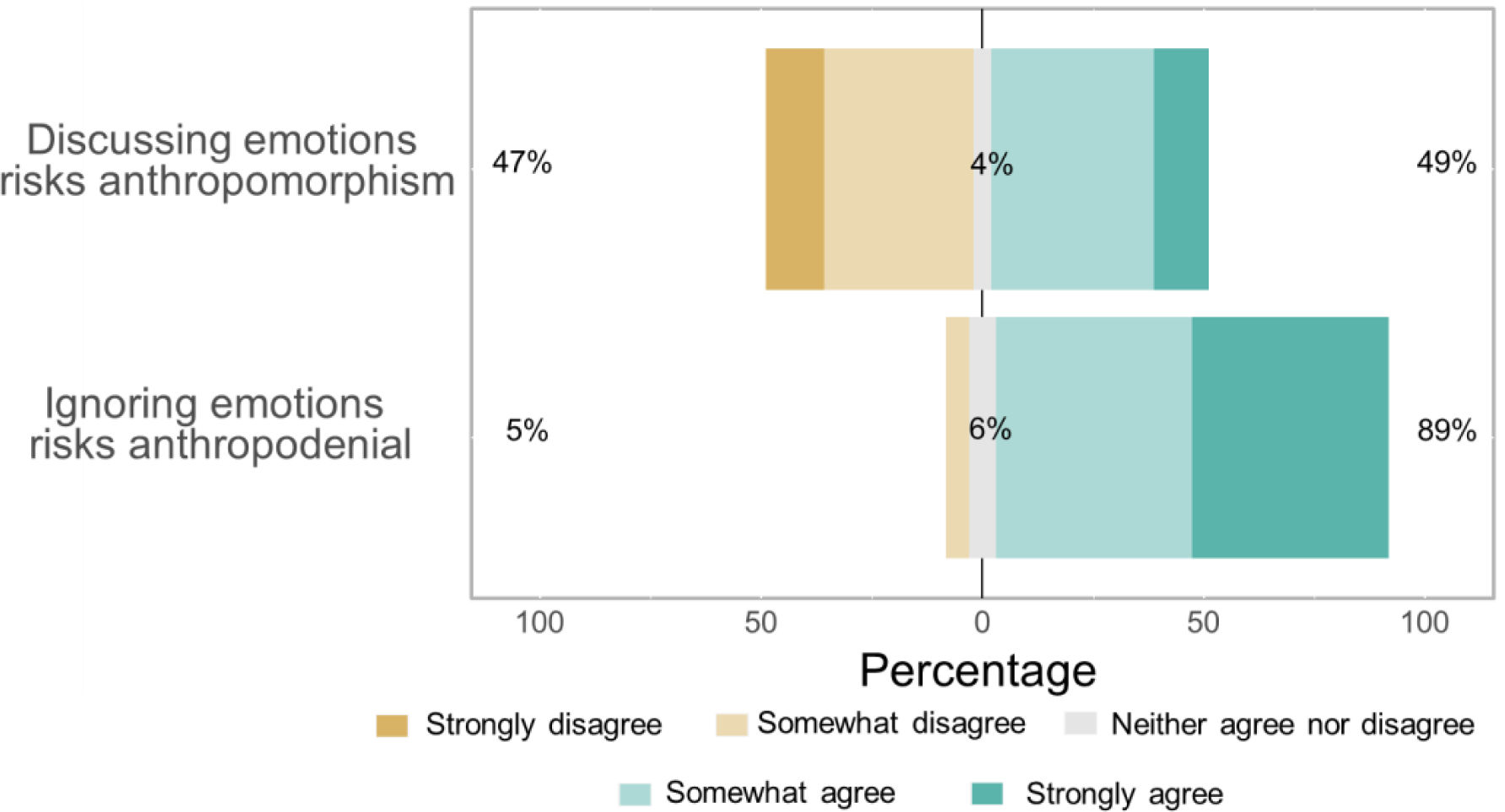
Respondents are divided on whether discussing animal emotions risks inappropriately anthropomorphizing animals’ experiences. Respondents generally agree, however, that ignoring the possible existence of animal emotions risks anthropodenial.

Among respondents who addressed both potential anthropic fallacies (n = 98), 39% agreed that both anthropomorphism and anthropodenial pose a risk. Another 46% agreed that anthropodenial is a risk, but disagreed that anthropomorphism is. Only 5% felt the opposite, that anthropomorphism is a risk but not anthropodenial (the remaining 10% selected “neither agree nor disagree for at least one statement). Thus, overall, nearly all researchers identify animal emotions as an area where one or more anthropic fallacies pose a risk to scientific progress, with a substantial minority (39%) identifying both issues as conflicting concerns.

### Perceptions of scientists’ abilities to measure emotions and consciousness in non-human animals

Based on the philosophical debate we outline in the introduction, it seemed possible that some respondents might argue that the question of whether a given animal has an emotional or conscious life might be untestable. To assess this possibility we asked respondents to state whether they believed that animals’ emotions and consciousness, separately, (1) could be measured using existing techniques, (2) could not be measured using existing techniques, but given reasonable advances in techniques would be measurable in the future, or (3) could not be measured using existing techniques, nor would it be possible to do so given reasonable advances in the future (see note in supplemental methods about selecting multiple options).

Only 11% of respondents said that animal emotions could not be measured now nor could they be reasonably expected to be measurable in the future. In contrast, 38% of respondents said that animal emotions can currently be measured with existing techniques, and another 52% said that although not currently possible, measuring animal emotions would become possible in the future given reasonable advances in techniques. Responses were essentially identical with regards to measuring consciousness (15% not possible now or in the future, 36% possible now, 49% not possible now but will be in the future).

### Perceptions of the field of animal behavior’s treatment of animal emotions

A large majority of respondents agreed that they would feel comfortable describing their beliefs regarding animal emotions and consciousness at a professional meeting (74%), while only 12% disagreed (the remainder neither agreed nor disagreed). Those that strongly agreed with the statement that they would be comfortable doing so were more confident in their average assessment of the presence or absence of emotions and consciousness in a given taxon (mean confidence rating = 3.6 for “strongly agree”, 3.0 for “somewhat agree”, 2.9 for other responses; “strongly agree” versus other categories p < 0.0002).

A large majority (71%) of respondents also agreed that the field of animal behavior “should do more to encourage research into non-human animal emotions, feelings, and consciousness” while only 12% disagreed. At the same time, only a slight majority (51%) of respondent agreed that the field of animal behavior already encourages researchers to consider animal emotions, while 35% disagreed.

### Animal Researchers’ Definitions of Emotion

As the final question of our survey, we asked respondents to define emotion. After reading respondents’ answers, two of us (MNZ and MFW) independently classified respondents’ answers as containing each of four components that many answers had in common: (1) reference to emotion resulting from internal and/or external stimuli, (2) reference to the function of emotion as motivating behaviors, (3) reference to subjective experience of an emotion/affective state or references to “mentality,” “consciousness,” “state(s) of mind,” or “feeling(s)”, (4) reference an explicitly social, communicative function of expressing internal state to others. These two independent assessments closely matched each other (>84% agreement). MNZ and MFW then collaborated to come to an agreement on responses that they had characterized differently.

81 respondents out of the 100 included in the broader survey provided a definition of emotions, perhaps indicating that for a substantial subset of respondents verbalizing their working definition was a challenge. Of the 81 definitions of emotion that respondents provided, 80 contained at least one of the components described above, with a slight majority (41/80) containing multiple components.

A majority of definitions contained a reference to emotions being a response to either internal or external stimuli (55%). A majority also contained a reference to emotions being subjective experiences or otherwise containing words related to consciousness or mindedness (56%). A substantial minority also identified emotions as functioning to motivate behaviors (40%). We note that earlier in our survey when we had asked respondents about their perceptions of the distribution of emotions, we reference “emotion that shape animals’ behavior.” It is possible that this framing shaped individuals’ likelihood of including this motivational aspect of emotion in their definitions, but given that this dimension of emotion was not the most common aspect of respondents’ definitions, any such unintended influence appears to have been limited in scope. A full breakdown of the distribution of these components and their overlap can be seen in Figure 5.

**Figure 5.**
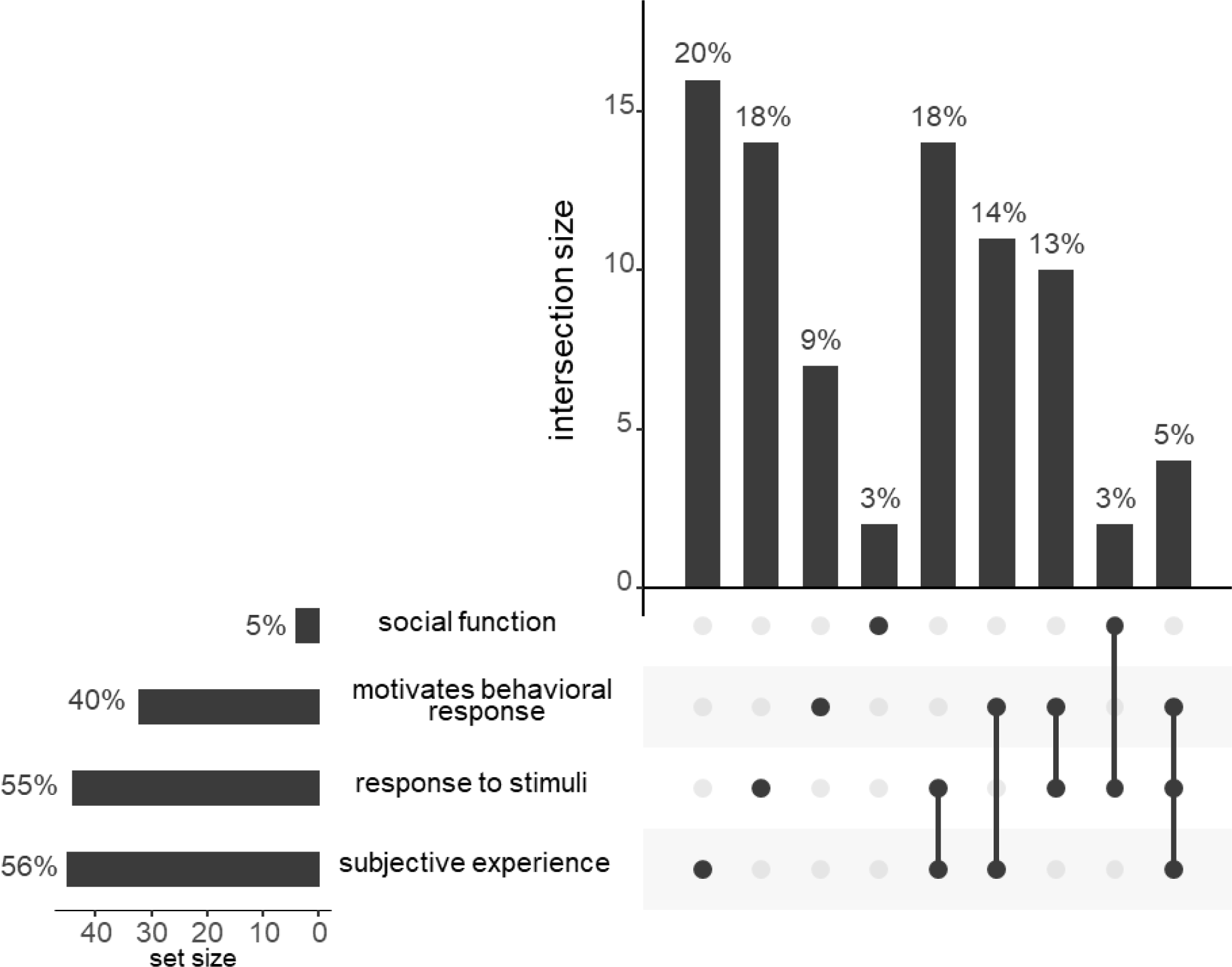
The distribution of components that respondents included in an open-ended request for their definition of emotions. Each vertical bar represents the percentage of respondents that included a given combination of potential elements that we scored in their definition. These components emerged organically after reading respondents’ definitions, rather than being pre-ordained. Horizontal bars represent the total percentage of respondents that included each individual element in their definition (rounding may result in apparent differences between horizontal bars and sums of vertical bars).

Despite substantial variation in the working definitions of emotions that respondents provided, individuals were no more or less likely to ascribe emotions to animals based on whether they included or excluded a given component. To assess this relationship while limiting the number of statistical tests performed, we first summed individuals’ Likert scale responses regarding their perceptions of the presence of animal emotions in each of the eight non-human taxa that we asked about (total possible range = 8-32). We then asked whether this total index value was predicted by whether an individual included each of the four components in their working definition of emotion. We found there to be no relationship between definitional components and overall assessment of the presence of emotions in non-human animals (t_73_; p > 0.3 in all cases, Figure S3). Within each of the eight individual taxa there was also no relationship between assessment of the distribution of emotion and individual definition components, after correcting for multiple hypothesis testing. In other words, although respondents may have had different models in their mind for what the word “emotion” conveys, this did not affect their assessment of the taxonomic distribution of emotions in animals.

## Discussion

In our survey, professional animal behavior researchers described a wide range of opinions regarding animal emotions. Animal behavior researchers ascribed both emotionality and consciousness to a broad swath of animal taxonomy and majorities identified questions regarding animal emotions as important and answerable. Several additional trends in researchers’ opinions emerge across a range of specific topics, including biases in researchers’ assessments, the relative risks of anthropomorphism and anthropodenial, and prospects for future progress.

### Taxonomic Distribution of Emotions and Consciousness

Animal behavior researchers ascribed emotions and consciousness to a wide range of animals, but consistently ascribed consciousness to a narrower range than they did emotionality. Respondents were less likely to ascribe consciousness than emotionality to “Most” or “All or Nearly All” members of all vertebrate taxa that respondents considered (non-human primates, other mammals, birds, fish, and reptiles and amphibians, Figure 1). This consistent pattern suggests that respondents perceive consciousness as a more derived trait than they do emotionality.

Respondents’ relative hesitancy to ascribe consciousness to animals in comparison to emotions is somewhat at odds with their stated assessment of the causal relationship between the two phenomena. Overall, 65% of respondents identified consciousness as being “somewhat necessary” or “essential” for animals to experience emotions. Based on this stated attitude, respondents should expect consciousness to generally evolve before emotions and that consciousness should be more widespread than emotionality. This apparent inconsistency in responses highlights the challenge that respondents, and indeed our fields, face when attempting to describe these phenomena and their relationships with precision. We purposefully did not define emotions for respondents but instead sought to quantify their unspoken sense of how emotions appear in non-human animals. Researchers’ possible struggle to apply a consistent standard in this regard is reflected in latent biases that our survey uncovered and in the varied definitions of emotions that they provided (see below).

### The Role of Phylogeny in Respondents’ Assessments

Researchers were more likely to attribute both emotional experiences and consciousness to animals that fell phylogenetically closer to humans, with their confidence in these ratings following the same trend (Figures 1-2). Large majorities ascribe emotions to most non-human primates, other mammals, birds, and cephalopods (in that descending order). This consistent phylogenetic trend is particularly striking given that a large majority (81%) of researchers stated that being phylogenetically close to humans was “not at all necessary” for animals to experience emotions.

Respondents’ assessment of cephalopod emotionality and consciousness provided the single exception to this phylogenetic pattern. Substantial majorities of respondents identified most “octopus, squids, and cuttlefish,” as showing both emotions (73%) and consciousness (65%), in contrast to insects and other invertebrates. Only 20-25% of researchers identify most invertebrates as displaying emotions or consciousness.

What is the source of this difference in researchers’ perceptions regarding cephalopods? Almost none of our respondents studied these species directly (2%). It therefore seems unlikely that this striking deviation in assessments is primarily the result of personal experience. What is more likely is that researchers have been exposed to the cognitive abilities of cephalopods through both academic and popular literature and films on the topic (e.g. *Other Minds* (30) and *My Octopus Teacher* (31)). The descriptions of individual behavior in these works may be so striking as to make a lasting impact on researchers’ assessment of the emotionality and consciousness of these animals. This possibility highlights both the powerful opportunity of charismatic portrayals of behavior to inform scientists’ understanding of systems with which they lack direct experience, but also the potential danger of such charismatic portrayals should they lack scientific rigor (we make no such assumption here).

### Additional Biases in Respondents’ Assessments

Respondents described many factors that had influenced their assessment of the presence of emotion in a given taxon while also stating that those same factors were not at all necessary for an animal to experience emotions. Many of these biases appear to result from the same subconscious heuristic: researchers are most likely to identify animals as showing emotions and consciousness if they share characteristics with humans. These characteristics included facial expressions, language, advanced cognitive abilities, and group living. But they also extended to characteristics not obviously shared by humans, including domestication (but see (32) for discussion of human self-domestication).

One of the biases that may have the greatest impact on the overall discourse regarding animal emotions and consciousness is a conservative bias in the absence of evidence. When researchers felt less certain regarding the distribution of emotions and consciousness in a given taxon, they were less likely to ascribe these phenomena to that taxon. Indeed, very few respondents were confident that a taxon lacked emotions or consciousness. In the absence of confidence that such phenomena exist, scientists assume that they do not. This attitude was summed up succinctly by one respondent who had stated that no members of any non-human taxa displayed consciousness. To explain this choice in a comment the respondent wrote, “As far as we know. I’m a scientist, I don’t guess.”

### Implications of Conservative Biases for Publication and Discourse

How might the overall conservative bias that we observe shape the landscape of publications on the topic of animal emotions? Because we asked our respondents about their confidence levels, we can play out a counterfactual in which we had screened out any participants who were not confident in their assessment—responding with a confidence of 3 or lower to a given question. Such a scenario would seem like a reasonable proxy for the types of respondents that might broadcast their arguments or beliefs regarding animal emotions and consciousness in a more rigorous, non-anonymous medium, such as a scientific publication. If we had considered only the perspectives of confident scientists, we would have reached even more extreme conclusions than we did above regarding the distribution of emotions and consciousness. Such respondents ascribed emotions to ‘most’ or ‘all or nearly all’ taxa at substantially higher rates for nearly all taxa considered, including other mammals (96% for high confidence respondents vs 89% overall), birds (95% vs 78%), cephalopods (95% vs 73%), fish (81% vs 52%), amphibians and reptiles (85% vs 49%), insects (45% vs 25%), and other invertebrates (38% vs 21%).

For the questions that we asked here, our overall conclusions did not hinge on respondents’ confidence. Yet, it reveals what might otherwise be a hidden bias in the “consensus” scientific view that develops on any given topic—if those that are most confident differ strongly from the median member of a field, those confident (and presumably louder) voices will be very likely to have an overweight influence on discussion both within the scientific community and in more public spheres.

The nature of these stronger voices has the potential to change over time. For example, among the respondents to our survey, substantially more respondents identified anthropodenial as a risk (89%) than said the same of anthropomorphism (49%). While we lack direct survey comparison on the topic, this distribution of thought likely represents a dramatic shift from the 20th century, when an aversion to anthropomorphism was dominant among animal behavior researchers (5, 29). That being said, nearly half of researchers still stated that anthropomorphism was a concern in animal emotion research. How do we reconcile these two concerns? Future research should strive to bridge this gap to both encourage lowering of prior barriers to research in areas related to phenomena previously assumed to be ‘private’, including mindedness, emotionality, and subjective experience, while being cautious of the real perils of anthropomorphism.

### Ethical Implications for Animal Care and Use

Animal scientists face both a legal and ethical imperative to consider the magnitude of animal suffering that a given scientific study will cause. Yet, reasonable people can and do disagree strongly about how much weight to give animal suffering in this calculation. Below we explore some of the potential implications of our respondents’ perceptions of animal emotions for animal research ethics, broadly. We take no personal positions on these topics here, and may or may not agree with the consensus views that our respondents describe.

A recent high-profile study that generated substantial ethical disagreement in the animal behavior community highlights the importance of researchers’ perceptions of animal emotions for deciding whether an experiment should be performed. A study was published in 2022 in the Proceedings of the National Academy of Sciences that made use of monkeys who had been the subjects of a mother-offspring separation protocol (for a different experiment) to ask whether mothers whose infants had been removed would interact with a soft monkey-like toy in the absence of their infant (Livingstone 2022). The separation was not conducted by the author of the paper and would have occurred in the absence of her involvement—she was simply asking whether mothers in such a circumstance would form attachments to soft, inanimate toys. Since the article’s publication, more than 260 researchers have co-authored a letter (33) in which they argued that the original experiment created so much animal suffering that any publication that made use of its existence was “unethical” and “outdated.” The letter’s arguments center on the psychological distress that they claim accompanies a disruption of the mother-offspring bond in primates. The authors state, for example, that primates “mourn” the deaths of offspring—a word that inherently invokes a complex set of emotional and physical reactions to the death of a close social partner.

The results of our survey cannot lead to a resolution of this debate. However, they do indicate that very large majorities of animal behavior researchers identify “most” or “all or nearly all” non-human primates as experiencing both emotions (98%) and consciousness (92%). It is therefore appropriate for IACUC committees to include considerations of emotional suffering in their determination of whether an experiment may be performed, and for this consideration to receive increased weight for non-human primates.

The value of an institutional process that considers extreme arguments from all sides without being dictated by them is especially great in ethical areas where individual scientists disagree strongly. How then do respondents’ assessments of animal emotions and consciousness align with institutional animal care and use committee (IACUC) guidelines in the United States? Overall, consistent with IACUC guidelines, respondents made substantial distinctions between the distribution of emotions and consciousness in vertebrate and invertebrate animals. For each vertebrate taxa, at least 49% of respondents identified “most” or “all or nearly all” members of a taxon as experiencing emotions (at least 39% said the same of consciousness). In contrast, only 21-25% of respondents said the same of insects and “other invertebrates.”

Yet, inconsistent with IACUC guidelines, respondents’ perceptions of cephalopods differ strikingly from other invertebrates. Based on respondents’ assessment of emotions and consciousness in these animals, it would seem reasonable to regulate their use in a way more similar to other vertebrates than to invertebrates. Cephalopods are currently excluded from legal protections that vertebrate animals enjoy, and are therefore not covered by most institutions’ IACUC oversight. Still, a few academic and governmental institutions have chosen to include cephalopods under their oversight umbrella without being legally required to do so (e.g. NASA Flight, Friday Harbor Laboratory, see also guidance from the Association for Assessment and Accreditation of Laboratory Animal Care).

### Conclusions and Optimism for Future Progress

Animal behavior researchers that responded to our survey were optimistic regarding our ability to understand animal emotions and consciousness. Most respondents to our survey assessed understanding animal emotions as an important area of inquiry, with a large majority (71%) agreeing that animal behavior, as a field, should do more to encourage investigation of “animal emotions, feelings, and consciousness”. Very large majorities asserted that animal emotions (90%) and consciousness (85%) can either be assessed using existing techniques or will be able to be assessed in the future given reasonable advances in technology.

The sum of our results indicate that respondents perceive (1) emotions to be taxonomically widespread and important in animals’ lives, (2) measurable now or in the near future, and (3) worthy of increased attention and research. This overall sentiment stands in stark contrast to the behaviorist sentiment of a century ago that emotions and consciousness represent either illusory or at best unmeasurable aspects of an animal’s life.

This study highlights the power of interdisciplinary collaboration to gain an understanding of a field’s consensus view regarding controversial or seemingly imprecise topics. Faced with a similar problem— i.e., the profusion of different but overlapping operative definitions of “genes”—a team of philosophers and biologists set out to determine the extent to which researchers in genetics and genomics study the same referent or merely talk past each other (the “Representing Genes Project”; (18–20)). This study revealed a “consensus gene” concept employed by its participants. Additionally, the study investigated “differences in which strategies were favored by biologists from different backgrounds,” and “whether the choice of strategies changed for human versus animal disease, and for physiological versus psychological disease” (34). The animal emotion literature could benefit from a similar definitional consensus to determine to what extent the limited research being done across different fields is referring to the same processes.

Our survey has made some progress towards this goal, revealing that researchers’ diverse definitions of emotions frequently included three shared elements– emotions involve an animal’s (1) response to a stimulus as a result of their (2) subjective experience of that stimulus and (3) that this response motivates a change in an animal’s behavior. Encouragingly, although only 5% of respondents included all three of these common elements in their free-response definitions of emotions, inclusion of these different components did not predict respondents’ assessment of the distribution of emotions across taxa. We suggest that these common definitional components, which emerged from respondents’ collective definitions of emotions, should serve as the foundation for shared use of language to facilitate conversations across disciplines surrounding these topics.

We share the optimism expressed by the respondents to our survey that progress regarding the distribution, functions, and limitations of animal emotions and consciousness will continue over the coming decades. We hope that this initial survey of sentiments and perceptions will serve as a baseline to which we or other researchers can compare in the future as new evidence emerges and our understanding regarding these questions becomes more precise.

## Acknowledgments

We gratefully acknowledge our sources of funding that made this work possible. MNZ has been supported by a National Science Foundation postdoctoral fellowship in biology (award #2109636) and a Klarman postdoctoral research fellowship from Cornell University. MFW was supported by the National Institute of Health (NIH) IRACDA (K12 GM000680/GM/NIGMS). This work was inspired by a conversation as a part of The Animal Behavior Podcast, which is supported by The Animal Behavior Society. We are grateful to all the researchers who took time out of their busy schedules to participate in our survey.

## Supplemental Material

### Respondents who selected multiple answers regarding scientists’ abilities to measure emotions and consciousness

We asked respondents to state whether they believed that animals’ emotions and consciousness, separately, (1) could be measured using existing techniques, (2) could not be measured using existing techniques, but given reasonable advances in techniques would be measurable in the future, or (3) could not be measured using existing techniques, nor would it be possible to do so given reasonable advances in the future.

We imagined these three possibilities to be mutually exclusive, but when designing the survey we unintentionally allowed respondents to choose multiple options if they wished. Many (24-35%) of respondents did so. When categorizing these responses, we took a conservative approach, treating respondents as choosing the most negative of the responses that they provided.

The largest category of these multiple respondents for both emotions (26/33) and consciousness (15/23) chose both “Possible using existing techniques” and “Not possible using existing techniques, but would be possible given reasonable advances in research methods.” We believe that these respondents meant to communicate, approximately, that while some aspects of emotions consciousness in non-human animals can be explored with existing techniques, getting at the core of what emotions and consciousness mean in other animals is a task that is not yet possible but will become so. In line with the conservative assumption we describe above, we treated these responses as most similar to “Not possible using existing techniques, but would be possible in given reasonable advances in research methods.”

All other multiple responses included the answer indicating that measuring emotions or consciousness was neither possible now nor would it be in the future given reasonable advances, so we interpreted them as intending to communicate this impossibility.

**Figure S1.**
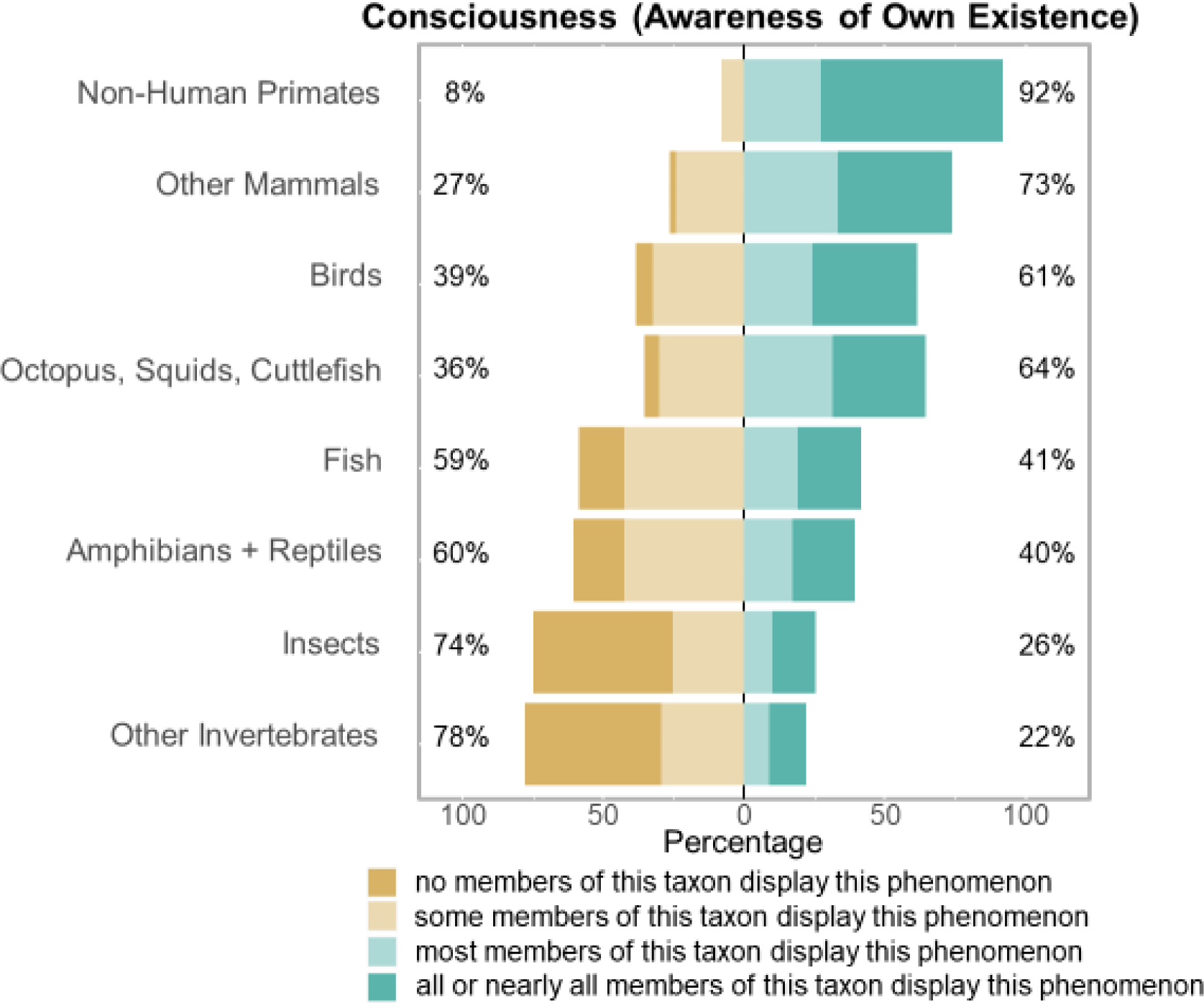
Respondents’ assessment of the distribution of consciousness across a range of non-human taxa.

**Figure S2.**
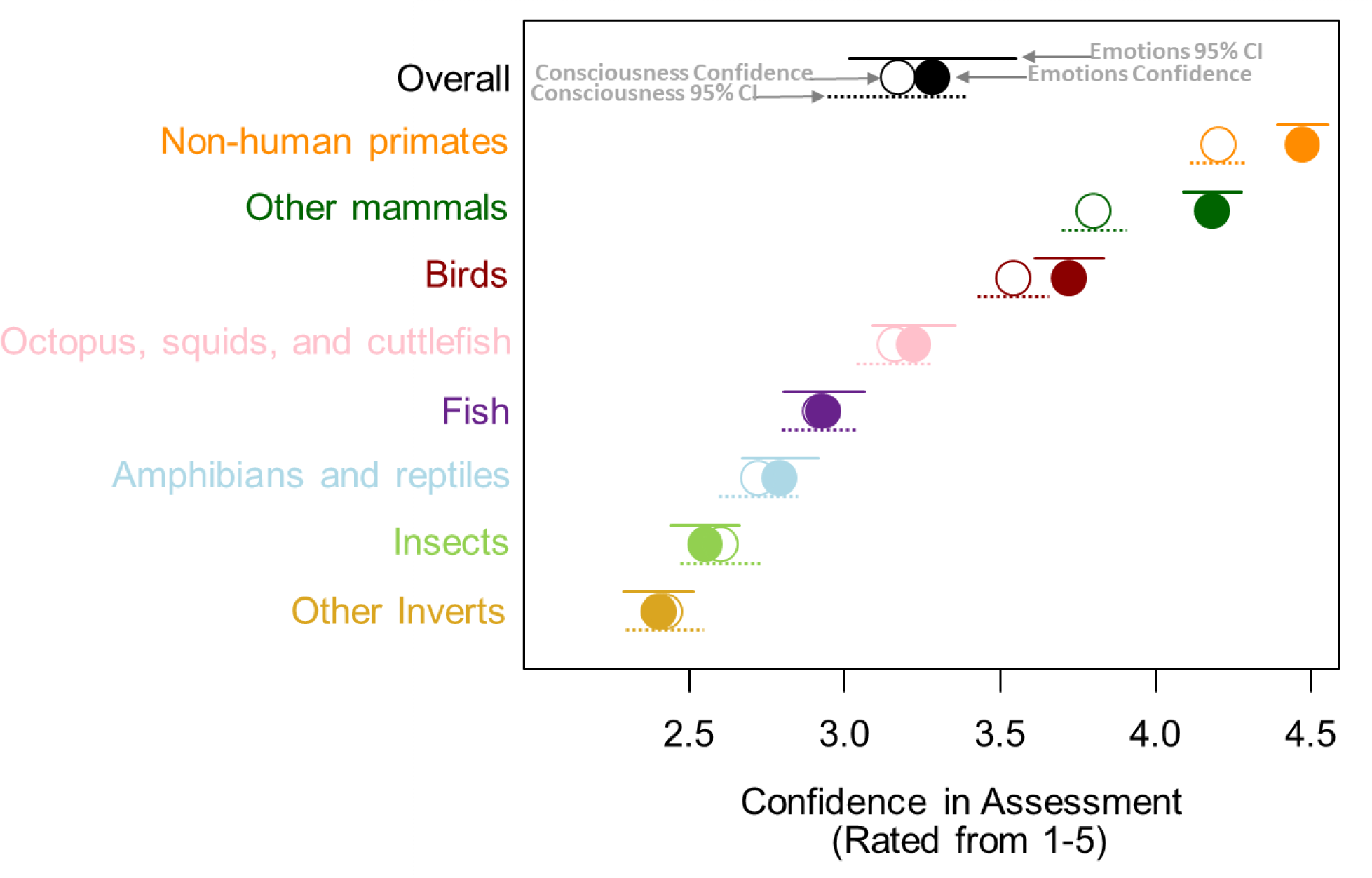
Respondents’ confidence in their assessment of the distribution of emotions (filled in circles) and consciousness (empty circles) in a given taxon were quite similar and showed the same taxonomic ordering.

**Figure S3.**
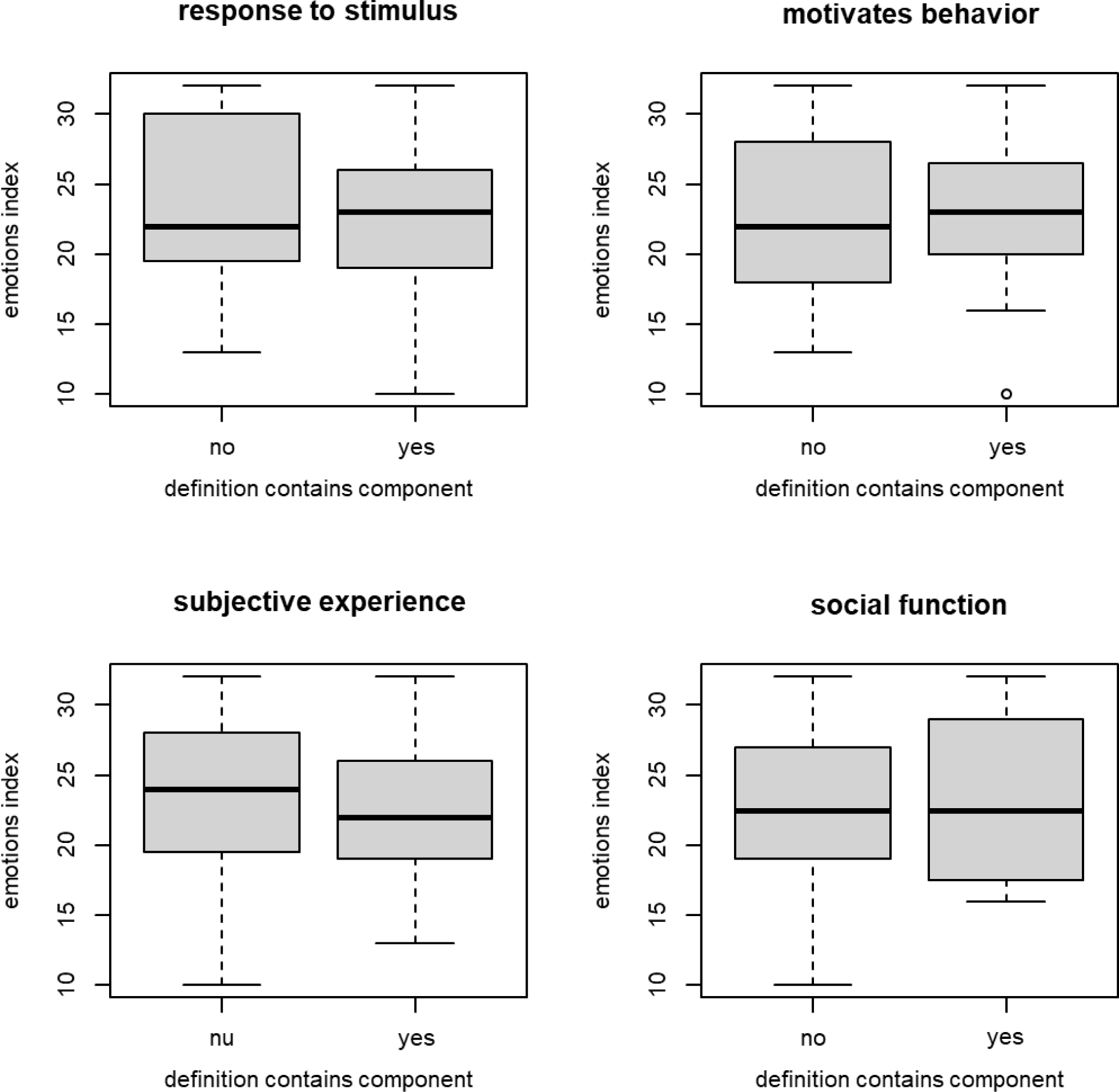
Respondents’ assessment of the distribution of animal emotions was unaffected by whether their working definition of emotion contained each of four different components common in individuals’ responses.

### Full Sample Survey

The following is a blank version of the survey that participants completed.

## Emotions and Animal Behavior

Q1 What is your institutional email address?

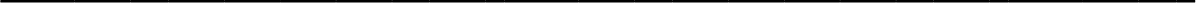

Q2 Which of the following best describes your career stage?

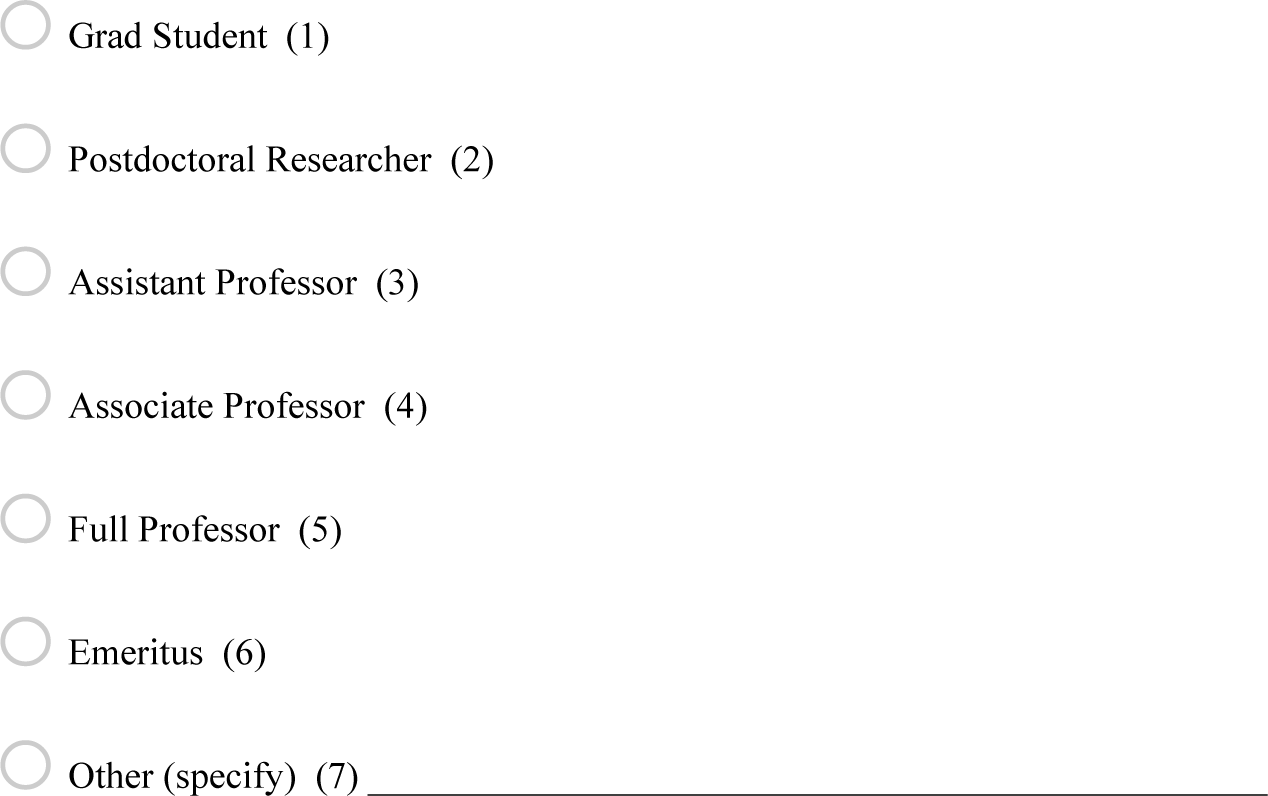

Q3 To which of the following disciplines do you belong? (choose all that apply)

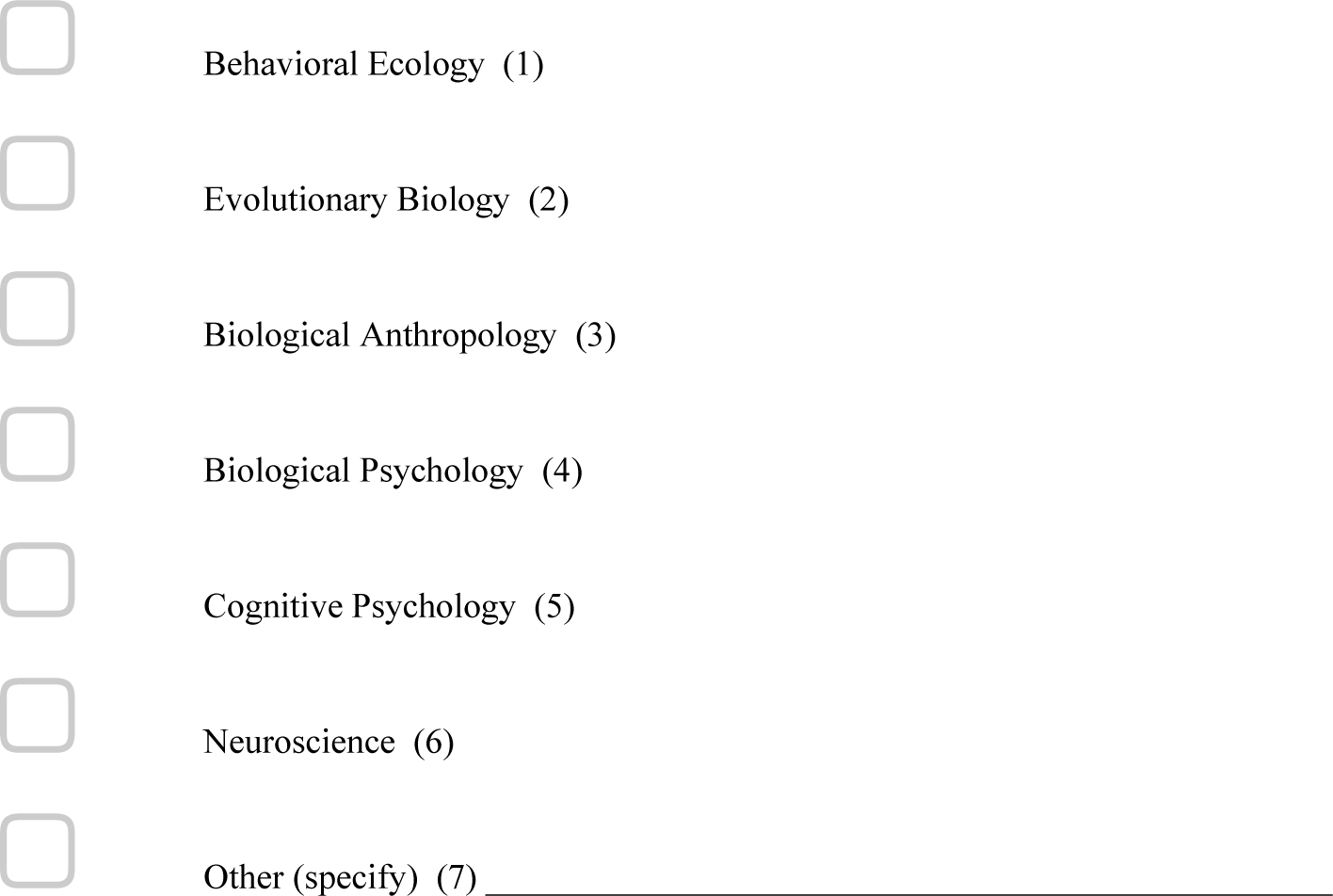

Q4 Which of the following taxa have you studied in the past 10 years? (choose all that apply)

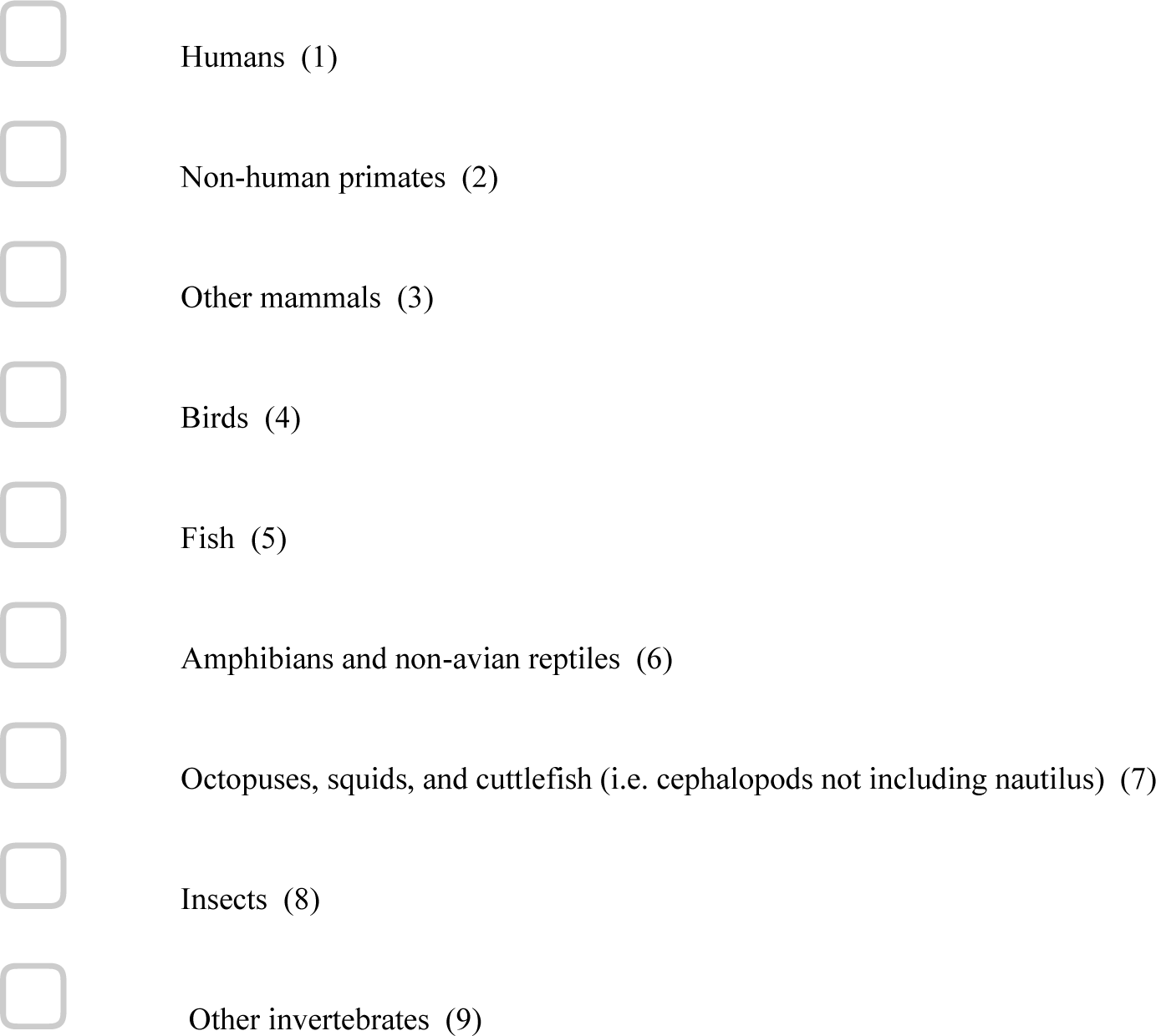

Q5 Which of the following pets or domesticated animals do you own or interact with daily? (choose all that apply)

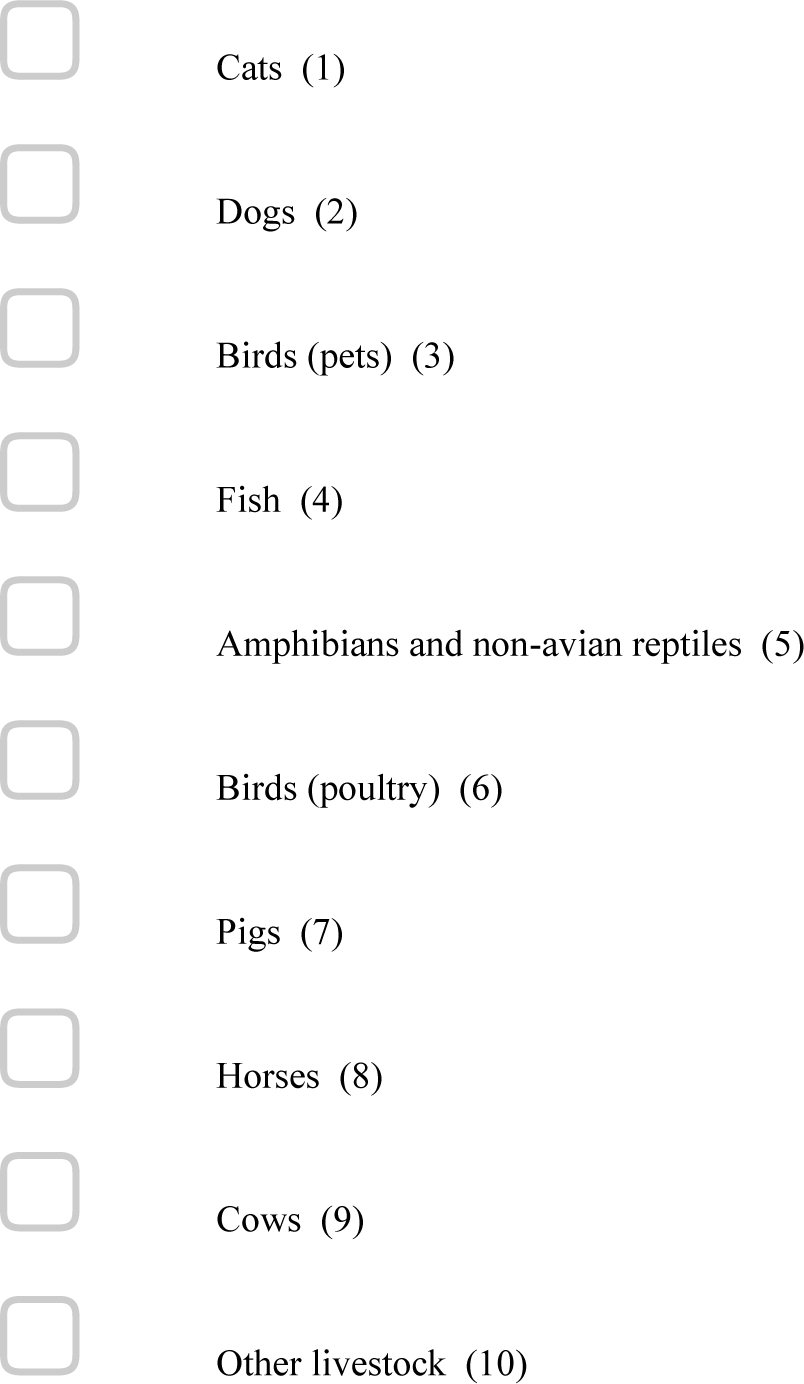

Q6 Some animals display emotional responses that in turn shape their behavior. Within the following taxa, how widespread do you believe this phenomenon to be? Please rate your confidence in your assessment from 1-5 with 1 being not at all confident and 5 being certain.

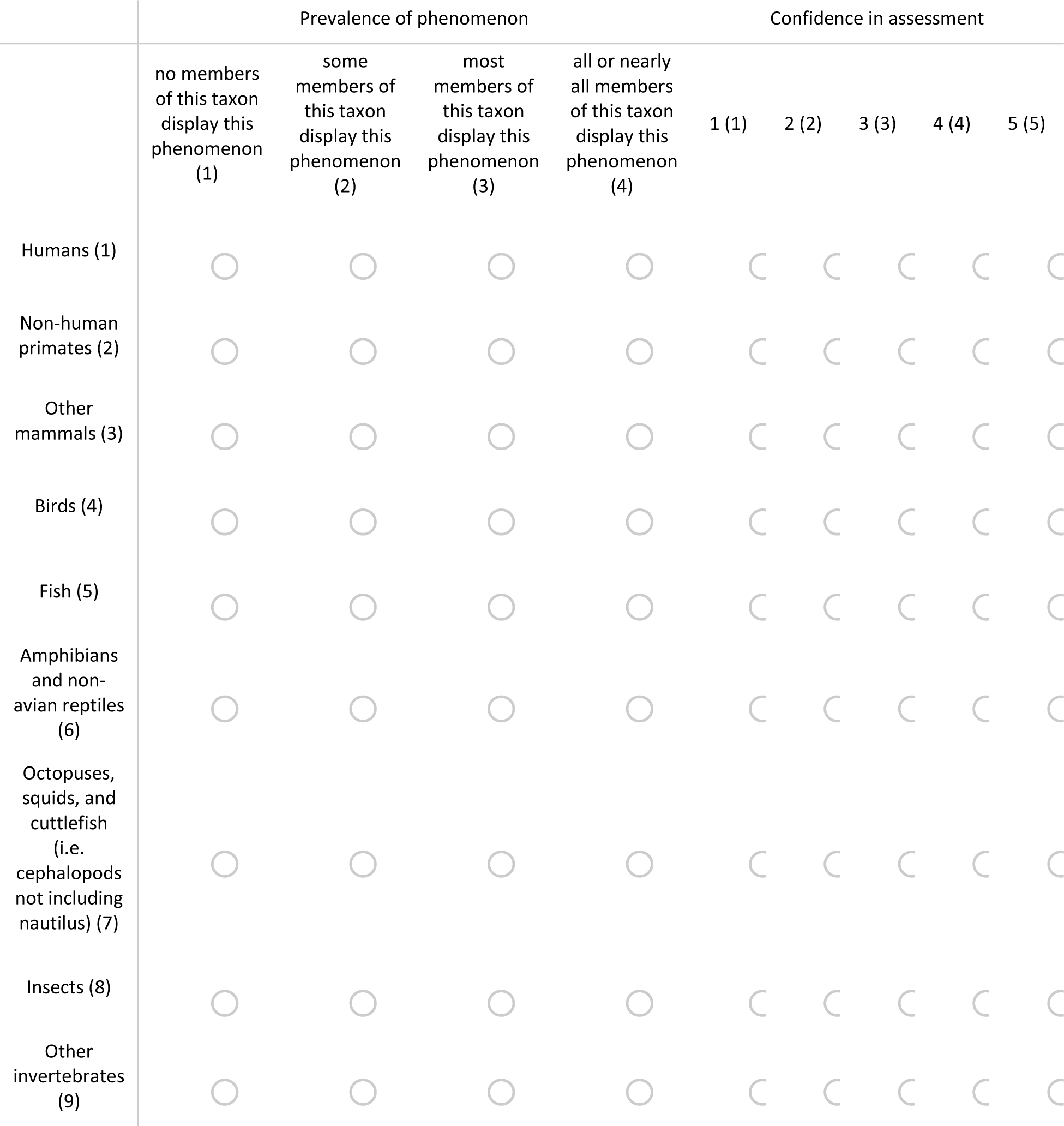

Q7 Some animals display consciousness, meaning that they are aware of their own existence. Within the following taxa, how widespread do you believe this phenomenon to be? Please rate your confidence in your assessment from 1-5 with 1 being not at all confident and 5 being certain.

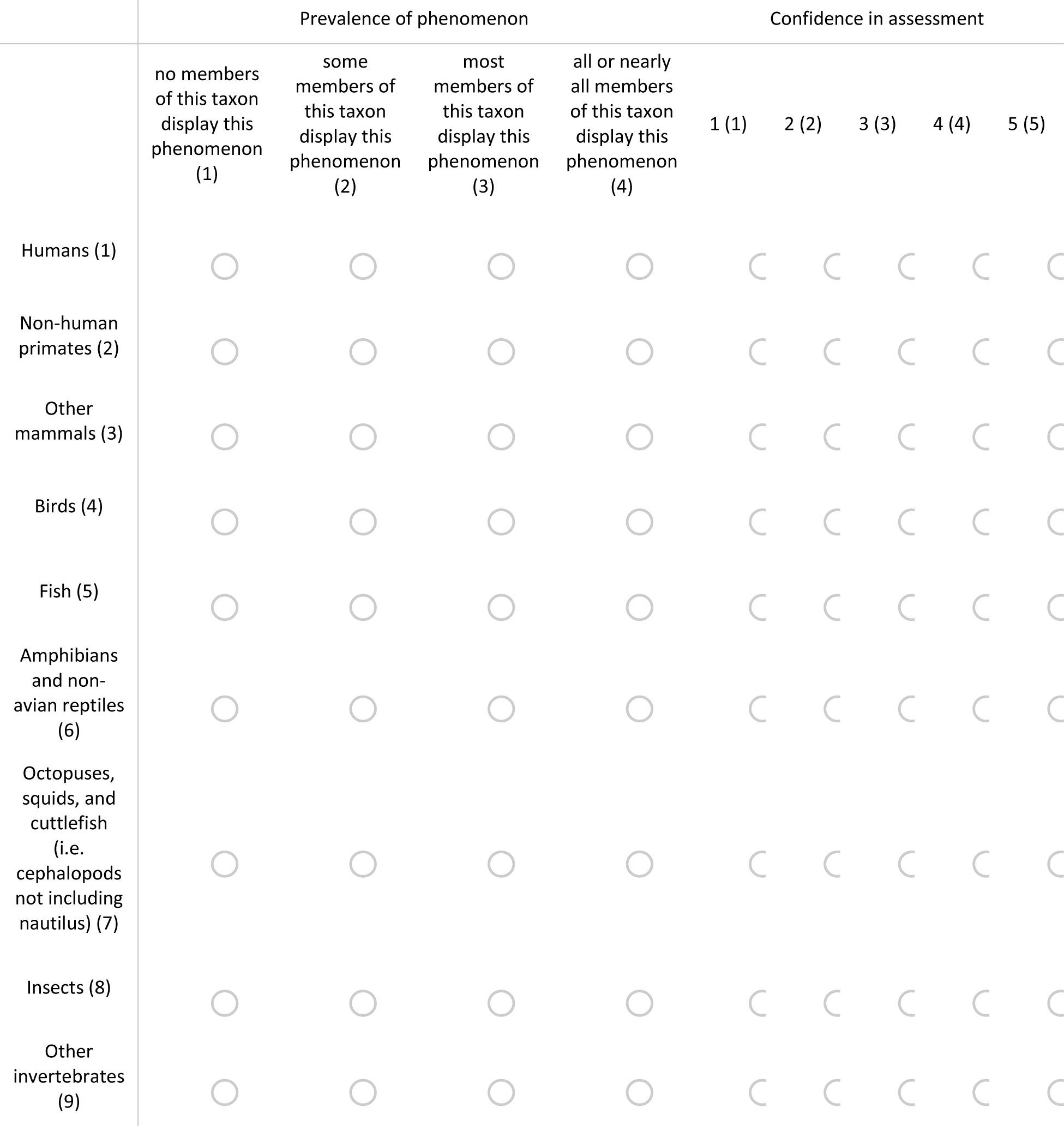

Q8 Thinking about your previous answers, how important were the following in whether you ascribed emotional responses to a given species?

|  | Not at all important (1) | Somewhat important (2) | Very important (3) |
| --- | --- | --- | --- |
| Phylogenetic closeness to humans (1) | ☐ | ☐ | ☐ |
| Consciousness (2) | ☐ | ☐ | ☐ |
| High level of sociality<br>(Engaging in individually differentiated social relationships) (3) | ☐ | ☐ | ☐ |
| Group living (4) | ☐ | ☐ | ☐ |
| Aquatic living (5) | ☐ | ☐ | ☐ |
| Advanced cognitive abilities (6) | ☐ | ☐ | ☐ |
| Use of language (7) | ☐ | ☐ | ☐ |
| Facial expressions that are observable to humans (8) | ☐ | ☐ | ☐ |
| Domestication/co-evolution with humans (9) | ☐ | ☐ | ☐ |

Q9 In your opinion, how necessary are the following traits for an animal to experience emotions?

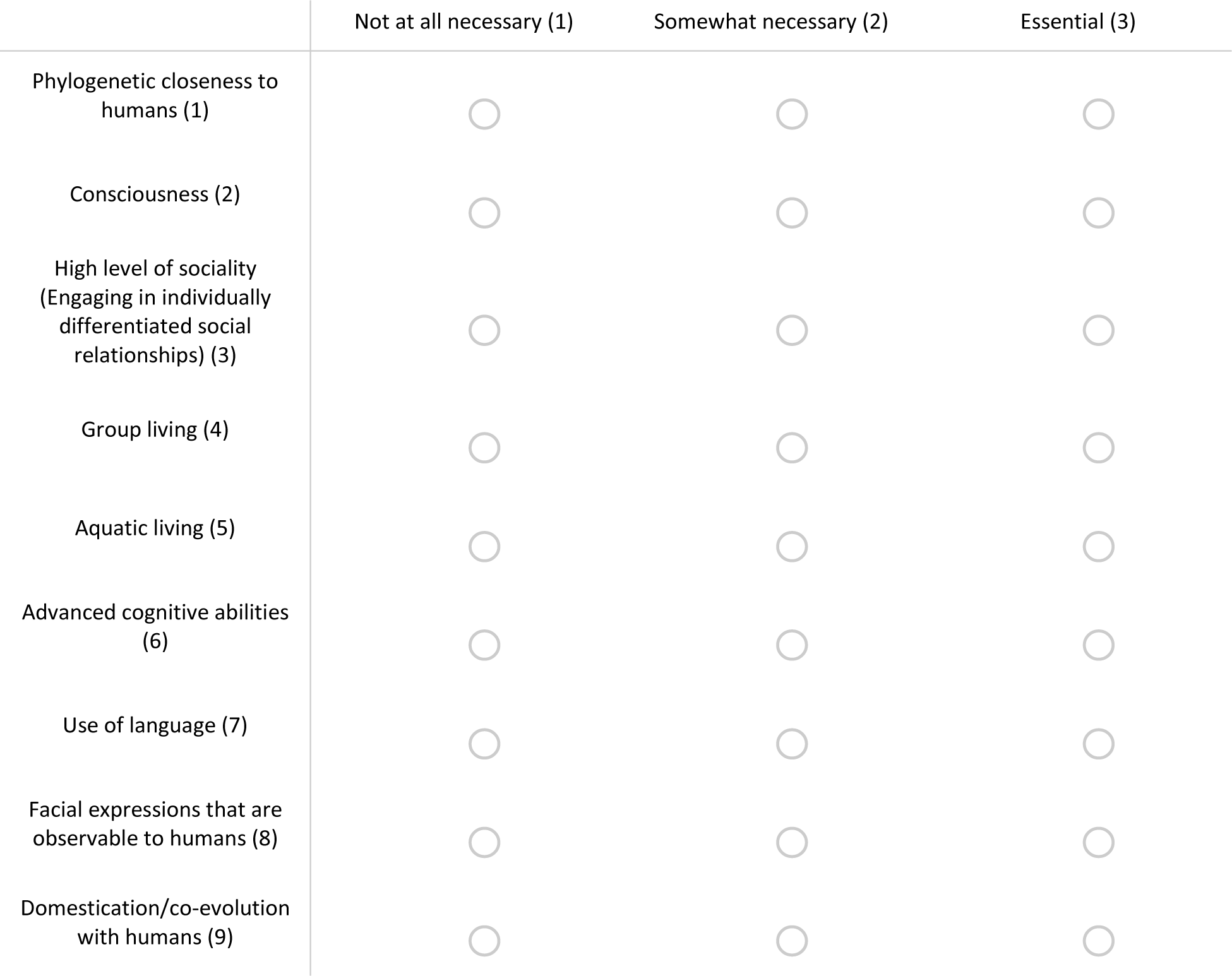

|  | Not at all necessary (1) | Somewhat necessary (2) | Essential (3) |
| --- | --- | --- | --- |
| Phylogenetic closeness to humans (1) | ☐ | ☐ | ☐ |
| Consciousness (2) | ☐ | ☐ | ☐ |
| High level of sociality<br>(Engaging in individually differentiated social relationships) (3) | ☐ | ☐ | ☐ |
| Group living (4) | ☐ | ☐ | ☐ |
| Aquatic living (5) | ☐ | ☐ | ☐ |
| Advanced cognitive abilities (6) | ☐ | ☐ | ☐ |
| Use of language (7) | ☐ | ☐ | ☐ |
| Facial expressions that are observable to humans (8) | ☐ | ☐ | ☐ |
| Domestication/co-evolution with humans (9) | ☐ | ☐ | ☐ |

Q10 To what extent do you agree with each of the following statements?

|  | Strongly disagree (1) | Somewhat disagree (2) | Neither agree nor disagree (3) | Somewhat agree (4) | Strongly agree (5) |
| --- | --- | --- | --- | --- | --- |
| Discussing the role of emotions in non-human animals' behavior risks inaccurately projecting human experience onto study subjects (i.e. anthropomorphization) (1) | ☐ | ☐ | ☐ | ☐ | ☐ |
| Failing to consider the role of emotions in non-human animals' behavior risks ignoring homologous mechanisms that are involved in both human and non-human behavior (i.e. anthropodenial) (2) | ☐ | ☐ | ☐ | ☐ | ☐ |
| Animal behavior, as a cross-disciplinary field, encourages researchers to consider the importance of emotions in shaping animals' behavior (3) | ☐ | ☐ | ☐ | ☐ | ☐ |
| Animal behavior, as a cross-disciplinary field, should do more to encourage research into non-human animal emotions, feelings, and consciousness (4) | ☐ | ☐ | ☐ | ☐ | ☐ |
| I would feel comfortable describing my beliefs regarding non-human animal emotions and consciousness with a colleague at a professional conference (5) | ☐ | ☐ | ☐ | ☐ | ☐ |

Q11 Measuring non-human animals’ emotional responses is (choose all that apply):

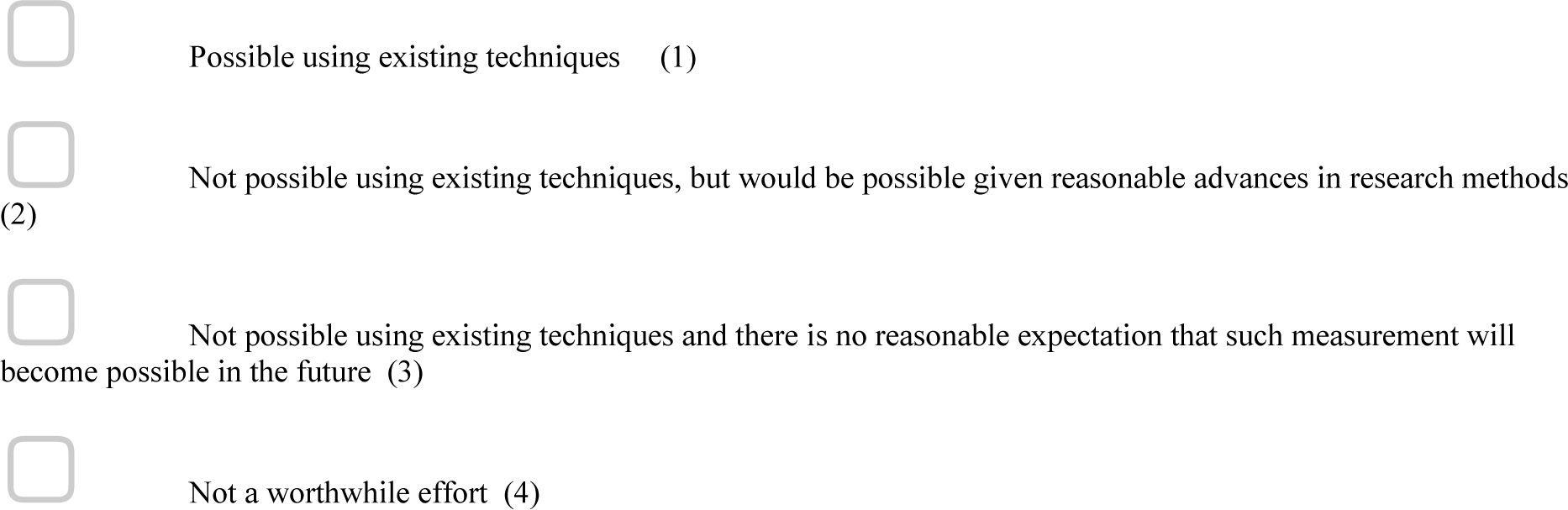

Q12 Measuring non-human animals’ consciousness is (choose all that apply):

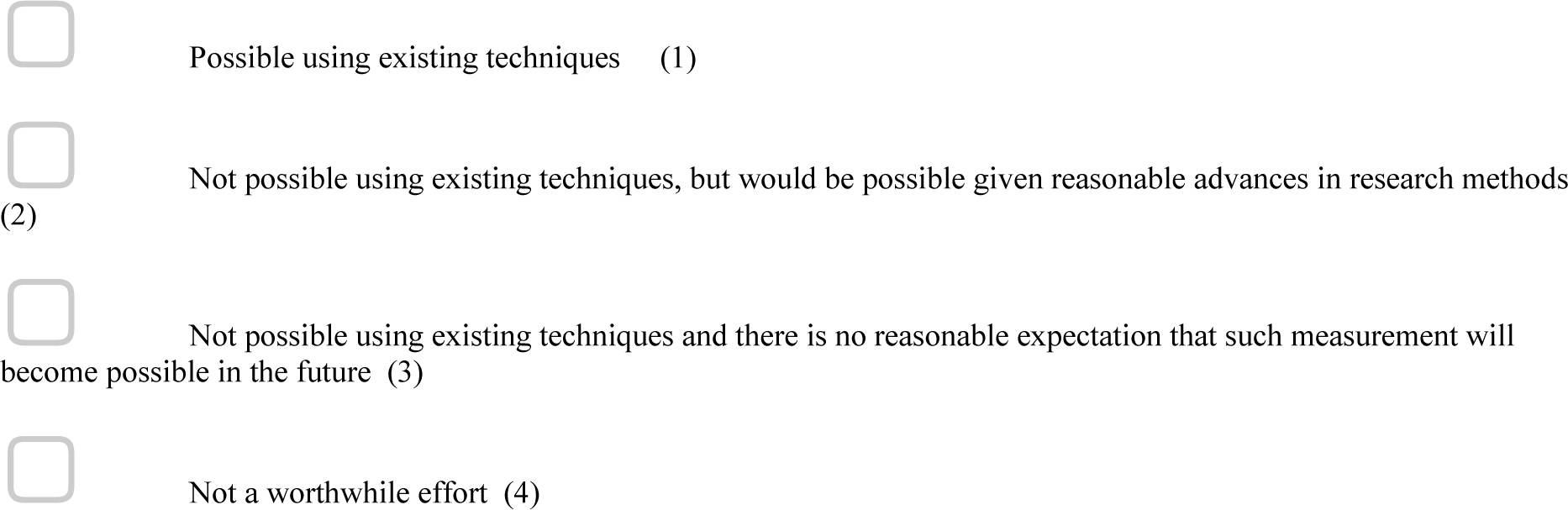

Q13 Please define emotion(s).

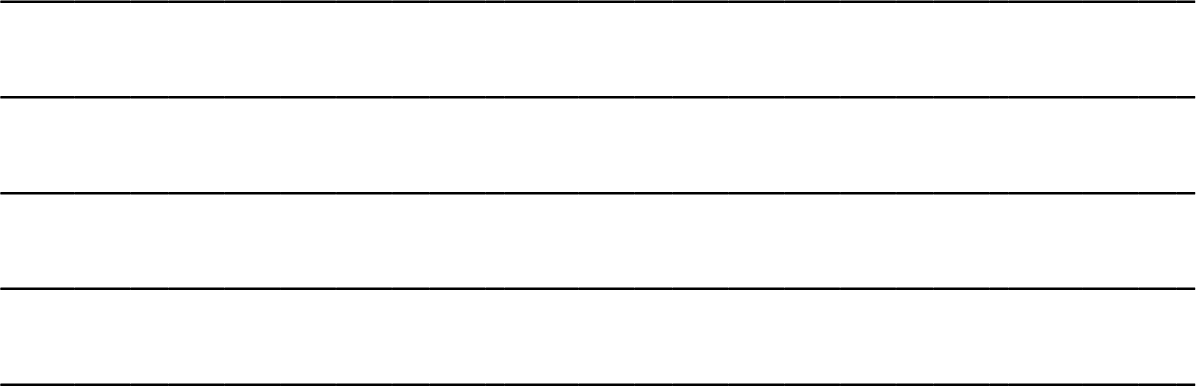

Q14 Do you or have you previously studied emotions or affective states in human or non-human animals?

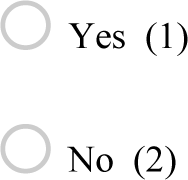

Q15 Do you differentiate between the terms “emotions” and “affect?

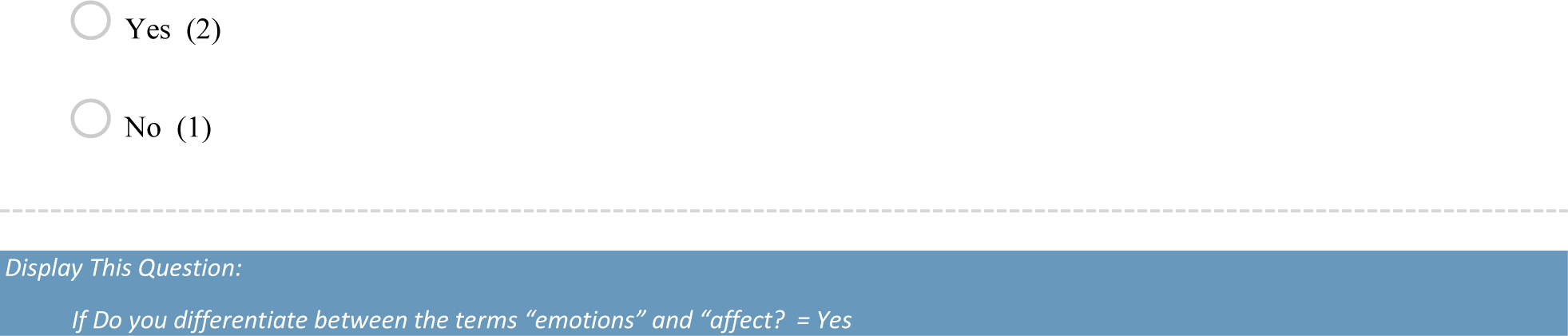

Q16

You have indicated that you differentiate between “emotions” and “affect.” Please reflect on how the differentiation between “emotions” and “affect” shaped the way in which you responded to the previous questions.

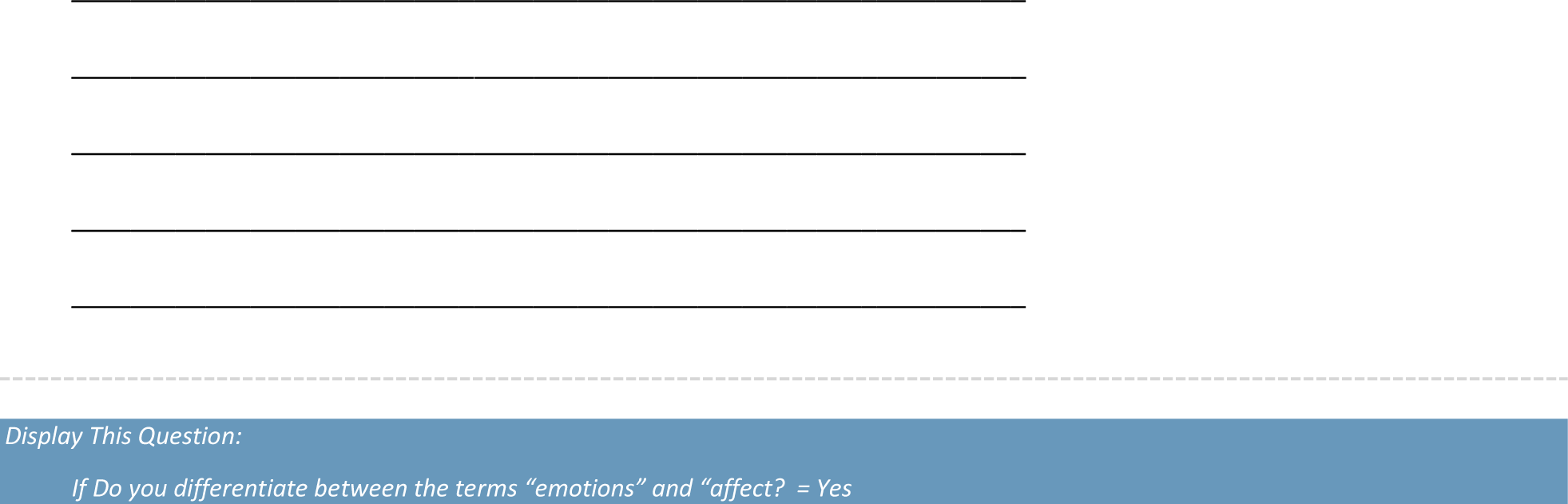

Q17

Some animals display affective responses that in turn shape their behavior. Within the following taxa, how widespread do you believe this phenomenon to be? Please rate your confidence in your assessment from 1-5 with 1 being not at all confident and 5 being certain.

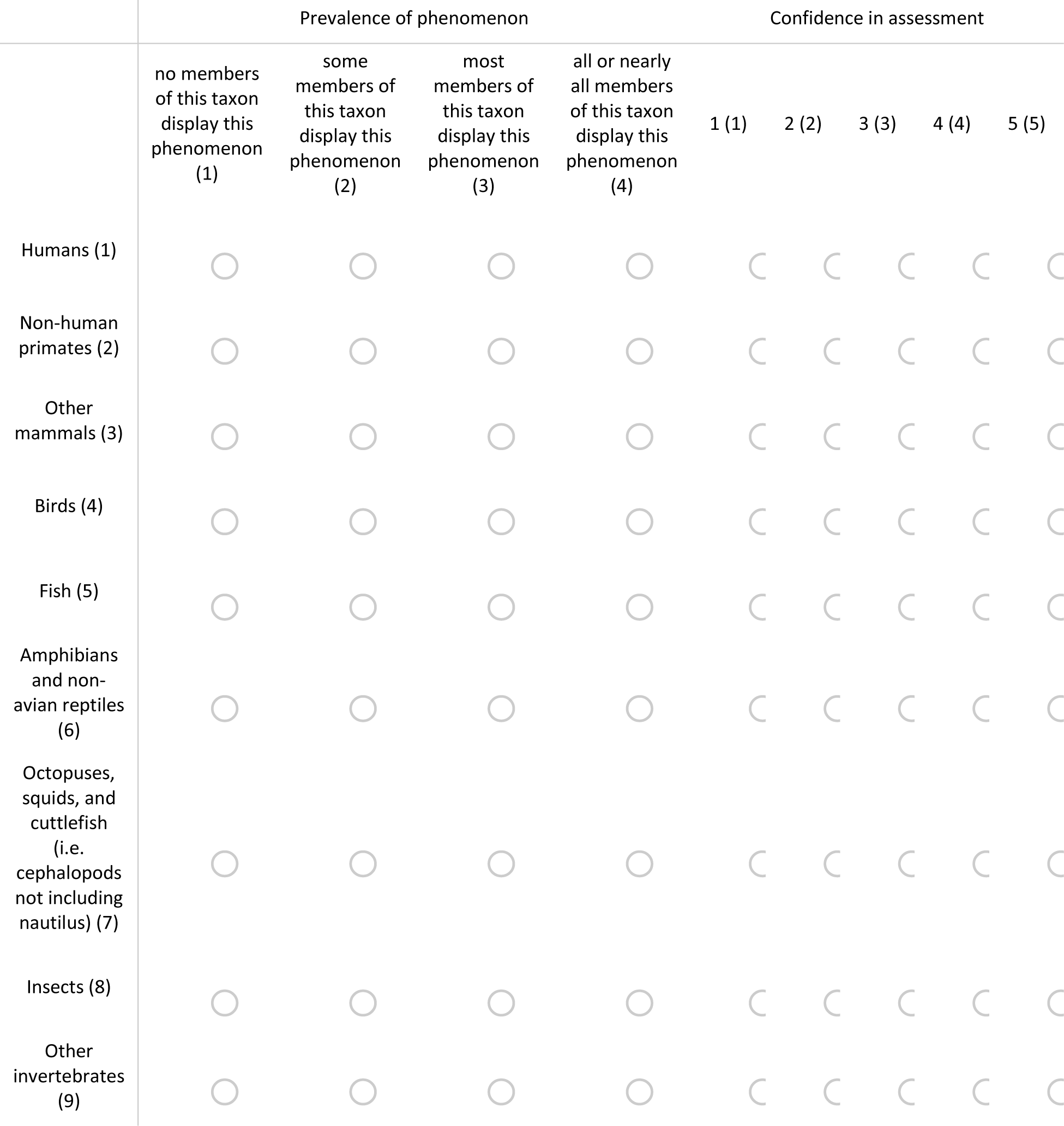

Q18 Please share any additional comments that you have on this topic or this survey.

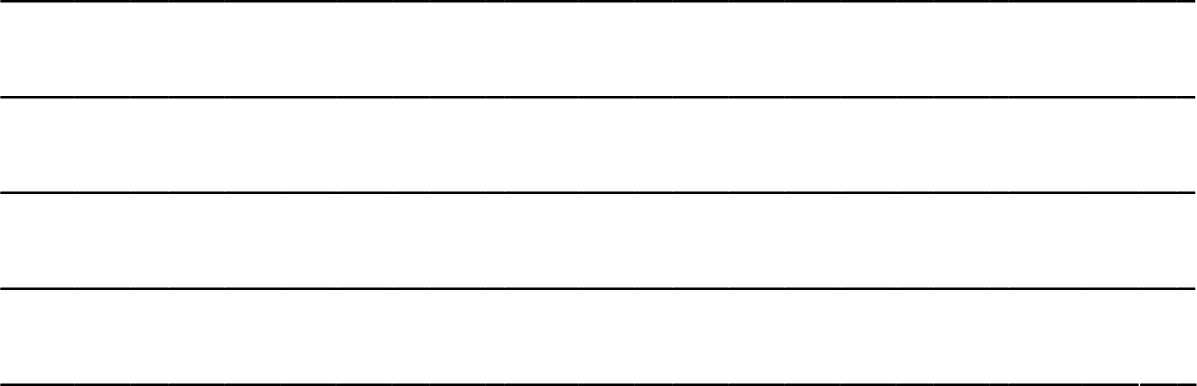

